# Genetic and environmental determinants of multicellular-like phenotypes in fission yeast

**DOI:** 10.1101/2023.12.15.571870

**Authors:** Bence Kӧvér, Céleste E. Cohen, Markus Ralser, Benjamin M. Heineike, Jürg Bähler

## Abstract

Multicellular fungi have repeatedly given rise to primarily unicellular yeast species. Some of these, including *Schizosaccharomyces pombe*, are able to revert to multicellular-like phenotypes (MLP). Our bioinformatic analysis of existing data suggested that, besides some regulatory proteins, most proteins involved in MLP formation are not functionally conserved between *S. pombe* and budding yeast. We developed high-throughput assays for two types of MLP in *S. pombe*: flocculation and surface adhesion, which correlated in minimal medium, suggesting a common mechanism. Using a library of 57 natural *S. pombe* isolates, we found MLP formation to widely vary across different nutrient and drug conditions. Next, in a segregant *S. pombe* library generated from an adhesive natural isolate and the standard laboratory strain, MLP formation correlated with expression levels of the transcription-factor gene *mbx2* and several flocculins. Quantitative trait locus mapping of MLP formation located a causal frameshift mutation in the *srb11* gene encoding cyclin C, a part of the Cdk8 kinase module (CKM) of the Mediator complex. Other CKM deletions also resulted in MLP formation, consistently through upregulation of *mbx2*, and only in minimal media. We screened a library of 3721 gene-deletion strains, uncovering additional genes involved in surface adhesion on minimal media. We identified 31 high-confidence hits, including 19 genes that have not been associated with MLPs in fission or budding yeast. Notably, deletion of *srb11*, unlike deletions of the 31 hits, did not compromise cell growth, which might explain its natural occurrence as a QTL for MLP formation.

## Introduction

Yeast species are defined as unicellular fungi and are found in the phylum Ascomycota. Within the Ascomycota, fission yeasts are found in the Taphromycotina sub-phylum, while budding yeast species belong to the Saccharomycotina (1). Phylogenetic data suggests that the last common ancestor of these branches, which existed around 500 MYA (1), was multicellular, but already possessed the ability to switch to planktonic growth (2). The primarily unicellular lifestyle of yeasts then repeatedly evolved by deploying a conserved set of transcription factors in each clade (2). Yet hundreds of millions of years later, yeasts still exhibit a range of multicellular-like phenotypes (3–9). Two widely used model organisms, the budding yeast *Saccharomyces cerevisiae* and the fission yeast *Schizosaccharomyces pombe,* exhibit flocculation (formation of multicellular aggregates) in liquid media and filamentous growth on agar plates (3–9). The latter is often coupled with the ability to invade agar (3–8). The fission yeast *Schizosaccharomyces japonicus*, closely related to *S. pombe*, also forms long filaments (10–12). Research on flocculation in *S. cerevisiae* has been driven by its role in brewing, where formation of flocs allows simple removal of biomass from each batch (9). Besides *S. cerevisiae*, *Candida albicans* is the yeast most studied for multicellular-like phenotypes such as flocculation (13), filamentous growth, and the clinically important phenotype of biofilm formation (14,15).

Importantly, these phenotypes are not *bona fide* multicellularity, in that they are temporary, lack committed cell fates, and the constituent cells replicate on their own rather than on a colony level (no germline). Nevertheless, filaments or clonal clumps (16) are made of genetically related cells, and even flocs might preferentially contain clonal cells (17). As such, these structures might constitute a level for natural selection to act on (18). Here we use the term multicellular-like phenotypes (MLPs) to refer to flocculation, surface adhesion, filamentous growth, invasive growth and biofilm formation, or any combination of these, in yeast species.

MLPs can give rise to emergent properties. Filamentation may facilitate foraging in nutrient-poor conditions (3–5,7,9). Similarly, flocculation could increase sedimentation in liquid media, thereby assisting the search for more nutrient-rich or less stressful environments (4). Alternatively, cell aggregates could share metabolic products of excreted enzymes as “public goods” and increase local nutrient concentrations in otherwise nutrient-poor environments (16,19). Indeed, at low sucrose concentrations, *S. cerevisiae* cells that clump together grow more efficiently than dispersed cells, primarily because they can share the products of the external sucrose invertase Suc2p, glucose and fructose (19). Moreover, inner cells of biofilms and flocs can be protected from environmental insults by the outer cell layers. Data from *S. cerevisiae* support a role of flocs in protection against high concentrations of ethanol, hydrogen peroxide, antifungals and UV (17). Moreover, surface adhesion and aggregation might protect colonies from being consumed by macroscopic predators, as has been shown for *S. cerevisiae* consumption by *Caenorhabditis elegans* (20).

Within a species, the regulators and effectors for different MLPs often overlap. The transcription factor Flo8p controls flocculation, filamentous growth and invasive growth in *S. cerevisiae* (21), and Mbx2 plays a similar role in *S. pombe*, although the two transcription factors are not orthologs (22). The cell-surface adhesion protein Flo11p is required for both invasive, and filamentous growth, and to some extent for flocculation, in *S. cerevisiae* (23). Similarly, the dominant flocculin Gsf2 is required for invasion, filamentous growth and flocculation in *S. pombe* (24,25). These observations, and other examples (8,13,20,26), suggest deep evolutionary and mechanistic connections between various MLPs and justify studying them together.

In *S. pombe,* sexual flocculation occurs between cells of opposite mating types as the first step of mating (27). Non-sexual flocculation occurs outside of mating, between clonal cells regardless of mating type (27) and depends on cell-surface flocculins which bind cell wall galactose residues in a Ca^2+^-dependent manner (24,28). *S. pombe* is also capable of forming filaments which can invade solid media (5,6,22,29). To quantify filamentation and invasion in *S. pombe,* assays have been developed to quantify the ability of cells grown on agar plates to resist washing (5,6,22,24,25,29–31).

During non-sexual flocculation, cell adhesion in *S. pombe* is primarily mediated by the flocculins Gsf2 and Pfl2-9 (24,25). These flocculins are positively regulated by the transcription factors Mbx2 and Cbf12, and are repressed by Gsf1 and Cbf11 (22,25,26,28,32). Another important aspect of MLP formation is the control of cell separation after mitosis. The genes coding for enzymes participating in septum digestion are activated by the transcription factor Ace2 (33). Both the *ace2* gene and the Ace2 targets are positively and negatively regulated by the transcription factors Sep1 and Fkh2, respectively (34,35), and this pathway could contribute to filamentation (7). Nitrogen starvation can trigger filament formation, but this requires a carbon source such as glucose which activates the cAMP/PKA pathway (5). Asp1, a kinase producing the inositol-pyrophosphate IP8, is required for filamentation through the cAMP-pathway, and overproduction of IP8 leads to flocculation (36). Interestingly, IP8 signalling is also associated with *mbx2* upregulation (37). Moreover, high iron concentration triggers surface adhesion and invasion of growth media (29). Furthermore, deletion of members of the Mediator Cdk8 kinase module and of some ribosomal genes (38,39) cause flocculation, while deletion of genes involved in mitochondrial gene expression (30,40) cause both flocculation and filamentous growth.

Much of the research on MLPs has been developed in the *Saccharomycotina* clade with studies in the budding yeast *S. cerevisiae* and *C. albicans* (3,4,9,14,41), and less is known about MLPs in *S. pombe* (6). A deeper understanding of the mechanisms behind MLP formation in *S. pombe* could provide insights into the latent capacity of yeast species to revert to ancestral multicellular phenotypes. To this end, we first analyse existing data obtained from model organism databases and show that while *S. pombe* shares a few regulators of MLP formation with *S. cerevisiae* and *C. albicans*, downstream effector cell-adhesion proteins are mostly not conserved between the three species. These results suggest novel mechanisms for MLP formation in *S. pombe*. In our lab, we observed that the natural isolate JB759 flocculates and weakly adheres to glass flasks in minimal medium. Here we screen for MLP formation across 57 non-clonal natural isolates (42), and find it to vary widely between strains and nutrient conditions. To understand the genetic basis of MLP formation in JB759, we apply a quantitative genetic approach revealing that MLP formation correlates with the expression of *mbx2* and flocculin genes. Through QTL mapping, we identify a causal frameshift mutation in *srb11*, functioning in the Cdk8 kinase module (CKM) of the Mediator complex. Deletion of CKM subunits caused an increase in *mbx2* expression, and MLP formation in these deletion strains depended on *mbx2*. To identify additional factors involved in MLP formation, we screened a genome-wide deletion library (43,44), and a library of long intergenic non-coding RNA deletions (45). We validated 3 known and uncovered 28 new genes involved in surface adhesion on minimal media. Of these, 13 had no previous annotation to MLPs in the budding yeast model organisms *S. cerevisiae* or *C. albicans*, while being genetically conserved. Interestingly, none of the adhesive natural isolates possess a null-mutation in the genes we observed as hits in our screen. We conclude that a null-mutation in *srb11* provides better growth efficiency compared to these genes, likely explaining the occurrence of only the former as a natural QTL for MLP formation.

## Materials and Methods

### Yeast strains

#### Segregants

Clement-Ziza et al. (46) created a segregant library by mating Leupold’s lab strain 968 *h^90^* (or JB50) and the South African natural isolate Y0036 (or JB759). Together with the two parental strains, we assayed 54 segregants from this segregant library, out of which two strains were identified as identical to other strains in the library upon sequencing. Named Rxx, eg. R45.

#### Natural isolates

Jeffares et al. (42) collected 57 non-clonal wild strains which span the natural diversity of *S. pombe*. Named JBxxxx, eg. JB1207.

#### ncRNA deletion library

Rodriguez-Lopez et al. (45) created a library of null-mutants for 141 different long intergenic non-coding RNA (lincRNA), each with multiple biological and technical replicates. During the confirmation step of our deletion library screen, all replicates of *SPNCRNA.1234*Δ and *SPNCRNA.900*Δ were verified by PCR and gel.

#### Prototrophic deletion library

The Bioneer V5 auxotrophic deletion library (43) was backcrossed with a wild-type strain to produce a prototrophic deletion library as detailed in (44). We used a copy of this library from Maria Rodriguez-Lopez, with multiple replicates for certain strains. All Mediator gene deletions were verified using PCR and gel. During the confirmation step of our deletion library screen, we chose a random set of 16 genes out of which 14 were successfully verified with PCR and gel.

#### *mbx2* overexpression strain

Using PCR, we amplified the coding sequence of *mbx2*, which we then cloned into the plasmid pJR1-41XL (47) using simple restriction digest and sticky-end ligation. The plasmid was amplified in “Mix & Go!” *E. coli* cells (Zymo Research) and extracted using a Qiagen mini-prep kit. The plasmid was transformed into the leucine auxotroph strain JB21 using a standard lithium-acetate protocol (48). As the plasmid had an *nmt1* thiamine repressible promoter, the transformed strains were first grown on EMM + 15 μM thiamine media and then placed in EMM for the experiments.

#### *CRISPR*-edited strains

We created a deletion of *srb11* in the JB50 background, and a deletion of *mbx2* in a *srb11Δ::Kan* background taken from the prototrophic deletion library. For more details, see the section CRISPR-Cas9 gene-editing.

| *Engineered strain name* | *Background* | *Genotype* |
| --- | --- | --- |
| *mbx2* overexpression strain (JB1774) | JB21 (*leu1-32 h-*) | pJR1-41XL::*mbx2* |
| *srb11* CRISPR knockout strain (JB1785, JB1786) | JB50 (968 *h90*) | *mbx2*Δ(CRISPR) |
| *srb11* and *mbx2* double-knockout strain (JB1782, JB1783, JB1784) | *srb11*Δ::*Kan* from deletion library | *srb11*Δ::*Kan*<br>*mbx2*Δ(CRISPR) |

### Media compositions and growth conditions

| *Name* | *Ingredients (for solid media add 2% agar)* |
| --- | --- |
| Yeast extract medium with supplements (YES - Rich media) | Yeast extract + adenine, uracil, histidine, lysine, leucine + 30g glucose / l |
| Edinburgh minimal medium (EMM) | EMM-N (Formedium) + 5g NH<sub>4</sub>Cl / l |
| Phosphate starvation | EMM-P (Formedium) + 1.81g NaCl/ l |
| Nitrogen starvation | EMM-N (Formedium) + 0.05g NH<sub>4</sub>Cl / l |
| EMM+ade (for segregants) | EMM-N (Formedium) + 5g NH<sub>4</sub>Cl / l +0.1g ade / l |
| EMM+thiamine (15 μM) | EMM-N (Formedium) + 5g NH<sub>4</sub>Cl / l + 15 μM thiamine |
| LB media for growing *E. coli* | LB (Formedium) |

Compounds were added at the following concentrations to these media: caffeine-10mM, rapamycin-100ng/ml. These concentrations were adopted from (49). RoToR HDA (Singer Instruments) compatible plates were poured using 40ml of media. Strains were always grown at 32°C, shaking at 160rpm (liquid cultures in tubes or flasks) or 80rpm (96-well liquid cultures) in an infors HT Incutron incubator.

### Orthology analysis

Candidate genes involved in MLP formation were obtained from relevant Gene Ontology (GO) terms (Supplementary Table 1). Orthology relationships between genes in *S. pombe* and *S. cerevisiae* were obtained from PomBase (50), while orthology relationships between genes from *C. albicans* and *S. pombe*, and *C. albicans* and *S. cerevisiae* were obtained from the *Candida* Genome Database (CGD; (51)). Such orthology annotations, in the case of PomBase, are a result of multiple algorithms and manual curation (50,52). In the analysis, genes were grouped into orthogroups to avoid confusion when it comes to accounting for paralogous genes (if one species had 5 versions of a gene, and another had 2, it was still counted in a single orthogroup). To extend our results, we also repeated this analysis after including genes from the Fission Yeast Phenotype Ontology (FYPO) (53), and the phenotype database from *S. cerevisiae* Genome Database (SGD; (54)) and CGD. Since SGD and CGD do not have a phenotype ontology, we obtained the respective tables of phenotypes, which were then filtered using keywords (Supplementary Table 1). We observed that *S. pombe* cell-adhesion proteins are not annotated as orthologs to other proteins in *S. cerevisiae* and *C. albicans.* To see whether the budding yeast cell-adhesion genes also lack orthologs in *S. pombe*, we examined cell-adhesion genes from *S. cerevisiae* and *C. albicans,* namely the *FLO* (FLOcculation) and *ALS* (Agglutinin Like Sequence) genes respectively.

### Sequence-, and structure-based queries for protein conservation

Our analysis, based on model organism database annotations, uncovered proteins assumed to be unique to *S. pombe* (including cell-adhesion proteins) and also proteins conserved across *S. pombe*, *S. cerevisiae* and *C. albicans*. We set out to independently verify this analysis using quantitative metrics. As a measure of protein conservation between *S. pombe* and *S. cerevisiae* or *C. albicans*, we used BLAST-P (55) which compares protein sequences and Foldseek (56) which compares protein structures. First, protein sequences were fetched from Uniprot using the Uniprot IDs obtained from PomBase. BLAST-P queries were then submitted through the Python function NCBIWWW.qblast(), with the arguments: database=”nr”, expect=1000, entrez_query=“txid237561[ORGN] OR txid5476[ORGN] OR txid559292[ORGN] OR txid4932[ORGN]” and hitlist_size=1000. Alphafold-predicted protein structures were fetched from https://alphafold.ebi.ac.uk/. Foldseek queries were then submitted through the Foldseek API using the recommended command:

*curl -X POST -F q=@PATH_TO_FILE -F ’mode=3diaa’ -F ’database[]=afdb-swissprot’* https://search.foldseek.com/api/ticket.

Both methods returned a list of candidate orthologs ranked by alignment scores. For all 25 “unique” proteins (including 15 cell surface adhesion proteins) and a random set of 50 “conserved” proteins we obtained the hits with the highest BLAST-P and Foldseek scores (Supplementary table 2). Statistical significance of the difference in these scores between “unique” and “conserved” proteins was assessed using a Mann-Whitney U test, separately for *S. pombe* - *C. albicans* and *S. pombe* - *S. cerevisiae* comparisons.

### High-throughput assays and *yeastmlp*

We developed two 96-well format high-throughput assays and a complementary software package for quantifying cell adhesion and flocculation. Both assays take advantage of the RoToR HDA colony-pinning robot (Singer Instruments), which can pin out yeast on agar plates in a 96-well arrangement. After conducting each assay, the data was analysed using our “Yeast Multicellular-like Phenotype” analysis package *yeastmlp* (https://github.com/BKover99/yeastmlp). Before each assay, yeast were transferred to YES plates from -80 °C glycerol stocks and were grown for 3 days at 32°C.

Our adhesion assay is a high-throughput variant of the conventional washing assay widely used in yeast literature (31). After pinning onto YES solid media from freezer stocks and incubating for 3 days at 32C, yeast were temporarily suspended in a 96-well plate filled with 200μl EMM in each well. Yeast were then pinned to agar plates with desired media conditions using the “7×7 squares” program on the RoToR, which pins 49 yeast colonies in a square arrangement for all 96 strains on an agar plate. This arrangement was chosen because it prevented yeast colonies from being washed off as a single self-adhesive colony, and allowed proper adhesion to agar. Yeast were then grown for 4 days following which they were imaged on a flatbed scanner (Epson V800 Photo), washed with water (constant 35ml/sec flow rate, 1s for each 7×7 square), and imaged again.

During analysis, *yeastmlp* takes a folder of pre-, and post-wash images, a 96-well map of strains, and an example “filled-out” plate, as arguments and returns adhesion values for each strain. To accurately discriminate between each square of cells, our algorithm creates a 96-well raw layout based on the example “filled-out” plate where each square contains growing strains. This raw layout is then fitted to each image separately in the pre-wash folder. Because the layouts are freshly generated for each analyzed image, our method should be robust to images acquired using different scanners. Furthermore, the individual fitting of layouts to each image allows robust quantification even if images in the same folder are slightly dislocated compared to each other (e.g., images from different quadrants of a flatbed scanner). Our assumption was, however, that pairs of pre,- and post-wash images are not dislocated with respect to each other. Finally, the algorithm expects at least one negative control (empty square) on each plate to correct for background intensity.

Since cellular density allows less light to pass through, high cell density is represented by low pixel intensities. Therefore, our measure of cellular density was inverse pixel intensity. After segmentation with the fitted-layout, mean cell densities in each square were calculated. In rare cases, the scanner returned slightly higher density values for washed colonies than pre-washing colonies; therefore, we decided to rescale all values to between 0 and 1 by dividing with the maximum inverse pixel intensity on a given image (meaning the darkest pixel). This resulted in robust and comparable estimates of colony density before and after wash in MLP-forming strains. We used a negative control (empty square) as a measure of background, which we subtracted from each measurement. Following this, the ratio of cell densities after and before washing allowed us to determine the fraction of cells sticking to the agar plate. Importantly, cells grown at the edges of the plate were more adhesive and produced less reliable measurements; therefore, our strains of interest were moved to the middle 60 positions. When a strain exhibited a pre-wash normalised pixel intensity value of less than 0.1, it was considered not growing on the given plate, and was removed from downstream analysis. Strains that had a mean pre-wash normalised pixel intensity value of less than 0.1 were removed from downstream analysis entirely. Throughout the paper, example raw data of plates before-, and after washing are shown using the viridis colormap, which appears perceptually uniform to the human eye (57).

For the flocculation assay, we used the “Archive” program on the RoToR to seed yeast cells in a non-TC treated 96-well plate in 200μl liquid media in each well. Cells were grown for 2 days and were imaged on a Tecan Infinite M200 plate reader which allowed measurement of optical density in each well of the 96-well plate at up to 225 different locations.

During analysis, *yeastmlp* takes as arguments a folder of .csv files returned by the plate-reader, the square root of measurements per well (e.g., 15 in the case of 15×15 measurements), a 96-well map of strains, and the location of the negative control well. The algorithm first subtracts the mean OD600 of the negative control from each well as a control for background light absorption. Our measurement for flocculation was the coefficient of variation (CV or standard deviation/mean) of normalized optical density measurements in each well.

To validate our high-throughput flocculation assay, we also measured flocculation using a simpler filtering assay. For this, yeast were grown overnight in YES, diluted to OD_600_=0.1 and inoculated into tubes with 5ml EMM. After 48h of growth, cells were resuspended by gently flicking the tubes and the culture was poured through a 30μm filter into a 50ml tube “A”. Flocculating cells stuck in the filter were washed into a second tube “B” and were completely resuspended using 10mM EDTA and vortexing. After measuring the OD_600_ of the content of both tubes, the ratio of flocculating to non-flocculating cells was determined as OD(B)/(OD(A)+ OD(B)). The two assays for flocculation showed a significant correlation (P=5E-15, Supp Fig 1). Generally, we consider the filtering assay more robust, while the plate-reader assay allows for higher throughput.

### Microscopy

Cells were grown from single colonies on YES plates over two days at 32C from single colonies in EMM or YES. After resuspending by shaking, 20ul of cells were placed on a glass slide and covered with a coverslip. Cells were then imaged on a Zeiss Axio-Imager Z2 with a Hammamatsu OrcaFlash 4.0 Camera with ZenPro2.3 software under bright field illumination using 20x air, 40x oil and 100x oil objectives.

### RNA-seq data

Clement-Ziza et al. (46) performed RNA-seq on the segregant library growing in EMM. We obtained a raw count matrix for our unbiased search of correlations between gene expression and MLP formation (46). Before the correlation analysis, raw count data was normalised using DESeq normalisation (58). For all additional “omics” data sources, see Supplementary Table 3.

### Finding shared upregulated genes across CKM deletions

After splitting segregants by their *srb11* haplotype, we performed differential expression analysis on the RNA-seq dataset from Clement-Ziza et al. (46) using DESeq2 (59). For further analysis, we used the top-100 upregulated protein-coding transcripts, by filtering for genes with log2FC >0.5 and Benjamini-Hochberg adjusted P-value <0.05, and finding the entries with the top-100 lowest P-values. We compared this gene set against upregulated genes taken from the microarray dataset from Linder et al. (60). Given that this dataset only contained sample means for each gene in each genotype, but no P-values, we simply took the genes with the top-100 largest log2FC values for both the *med12Δ* and *srb10Δ* genotypes. The intersection of the 3 gene sets then identified 15 genes which are upregulated in *srb11* truncation and *srb10Δ* and *med12Δ* genotypes.

### Finding overlap between genes upregulated upon CKM deletion and Mbx2 targets

Once we identified genes upregulated across the CKM deletion strains, we set out to compare this set with known targets of Mbx2. Kwon et al. (25) have identified targets of Mbx2 by collecting microarray and ChIP-chip data. We considered genes to be upregulated in the microarray dataset with log2FC >1 and Bonferroni-adjusted P-values <0.05. Furthermore, as Kwon et al. (25) have already performed quality control and filtered the ChIP-chip data for likely targets, we used every gene in that dataset. By finding the intersection of these 3 gene sets we identified 5 genes, including *mbx2*, which are upregulated in CKM deletion strains, likely through the activity of Mbx2. That *mbx2* itself is part of the gene list reflects that it binds its own promoter to activate its gene.

### DNA sequencing data

Genotyping of 44 strains in the segregant library was done by Clement-Ziza et al. (46). Briefly, they performed whole genome sequencing on the two parental strains with high coverage, and after calling short variants, they inferred the genotypes of segregants at each locus using bulk RNA-seq data. There remained 10 strains in the library that were not analysed by Clement-Ziza et al. (46), likely because of their strong adhesive phenotype affecting downstream procedures.

We sequenced the remaining 10 strains in the library, and also the strain R4 as a control to compare our methods with that of Clement-Ziza et al. (46). DNA extraction was done using a protocol obtained from Daniel Jeffares (personal communication, 2022) which involves spheroblasting followed by lysis, RNA and protein removal, and DNA extraction with the Qiagen Genomic-tip (20/G) protocol. Briefly, a 20ml culture of cells was grown up in YES at 32°C and harvested by centrifugation (3000xg for 15 min at 4°C). Cell walls were digested using lysing enzymes of *Trichoderma harizanum* dissolved in 50mM citrate-phosphate buffer, pH 5.8, with 40mM EDTA and 1.2M Sorbitol, and incubated for 1.5h at 32C to generate spheroblasts. Cells were then centrifuged at 3000rcf for 10 min at 4C and supernatant was removed. Following this, the Qiagen Genomic-top (20/G) protocol was followed, from page 37, step 8. Finally, high-quality DNA was isolated using isopropanol and ethanol precipitation. Library preparation and Illumina NovaSeq paired-end sequencing at well above 100x coverage was done by Azenta.

The resulting FASTQ files were checked for quality using FASTQC (61), following which we performed adapter trimming using Cutadapt (62) using default parameters. The reads were mapped to the *S. pombe* reference genome using BWA MEM (63) using default parameters. The resulting alignments were then processed through the GATK short variant discovery pipeline (64) using default parameters. In this pipeline, we used Base Quality Score Recalibration based on the .vcf file listing all known variants discovered by Jeffares et al. (42). Although we collected haploid *S. pombe* samples, GATK assumed a diploid status during genotyping, which we used as a quality measure and discarded calls with heterozygous status, similarly to what was done before (46).

We also accessed the original FASTQ files from the 2 parental strains (ENA accessions: ERX007392, ERX007395), and together with our newly sequenced 11 strains (total of 13), we genotyped them to produce a variant call format file. This was then integrated with the genotype table from the supplementary material of Clement-Ziza et al. (46). Uncalled variants where both the preceding and subsequent variants came from the same haplotype were imputed for each strain.

To evaluate our genotype calling pipeline, we compared our calls for the parental strains versus the calls made by Clement-Ziza et al. (46). Using our pipeline on the sequencing data from (46) we saw that for JB50, our pipeline identified a wild-type genotype for 4475 out of 4481 (99.87%) variants found by Clement-Ziza et al. (46). For JB759, our pipeline identified the same alternative allele as Clement-Ziza et al. (46) 4418 times out of the 4481 (98.59%) variants. To compare our sequencing protocol and variant identification pipeline to that used in (46), we compared the calls for the segregant R4, for which we sequenced DNA and which had variants called based on RNA-sequencing data in (46). We found that the called SNPs differed at 26 loci out of 4481 (0.58%), and concluded that both our computational genotyping and DNA sequencing pipeline was robust.

During our quality control step, we identified an extremely high overlap in variant calls of the two segregants R4 and R45 (>99% overlap), and the segregant R48 and the parental strain JB50 (>99% overlap). We therefore renamed R45 as R4_45 and R48 as JB50_48 and omitted them from further strain specific analysis. Because we now had two high-coverage replicates for R4, named R4 and R4_45, two for JB50 including the original sequence from (46) and newly sequenced JB50_48, and high coverage for our other newly sequenced samples, we used these sequences to call further short variants previously not reported in (46). Our criteria was that these variants should be called as homozygous by GATK haplotype caller with different genotypes in the two parental strains, and that the variant should match between JB50_48 and the originally sequenced JB50 strain, as well as between R4 and R4_45. We called an additional 387 short variants, with an average length of 2.59 nucleotides for the newly called JB50 alleles and 2.88 nucleotides for the JB759 alleles. These stand in contrast with the variants called by Clement-Ziza et al. (46) which were on average 1.22 and 1.15 nucleotides in length, meaning that we mostly identified indels while the previously called variants were mostly SNPs. In the segregant strains genotyped only using RNA-seq by (46), these variants were imputed to match the haplotype of preceding and subsequent variants in the genome.

In the end, our sequencing efforts extended the dataset from 44 to 52 segregants, from 4481 to 4868 short variants, and from 685 to 812 haplotype blocks (Supplementary Table 4). Additionally, our raw sequencing data has been archived in the European Nucleotide Archive (www.ebi.ac.uk/ena/), with study accession PRJEB69522.

For the natural isolate library, we obtained genomic data from Jeffares et al. (42) in a processed variant call format.

### QTL-analysis

Quantitative trait loci analysis was done using the R-based RFQTL algorithm described in (46,65), downloaded from http://cellnet-sb.cecad.uni-koeln.de/resources/qtl-mapping/. This method is based on the Random Forest machine learning algorithm. Briefly, short variants (SVs) and phenotypes are used to build decision trees, objects in which SVs with the highest explanatory power partition the phenotype data through a hierarchy of steps. Random subsets of data used for each decision tree give rise to a so-called random forest. There is always a “competing” collection of SVs being simultaneously considered, rather than a single variant, as it is commonly the case for univariate statistical tests used for similar purposes (65). The hierarchical partitioning, and the simultaneous consideration of multiple variants help to account for epistatic mechanisms and achieve higher fidelity QTL hits (65). Statistical significance is obtained as follows: first, an importance score (“selection frequency”) is calculated for each SV on a small set of forests, and they are then compared to a null-distribution of importance scores coming from a large set of forests with bootstrapped data (65). For our QTL analysis, we generated the importance scores from 100 forests with 100 trees each and created the null distribution using 20,000 permutations of 100 forests with 100 trees each. The number of permutations was set such that P-values of genome-wide significance could be achieved given Bonferroni-correction for multiple testing:

n_permutations > n_haplotype_blocks / sig_threshold (= 803/0.05 = 16060).

### Identifying variants causing a premature stop codon or frameshift

Identification of premature stop codons and frameshifts was done using a bespoke Python script that takes the reference genome .fasta file and genome annotation .gff3 file from PomBase, and an input of our variants of interest. After QTL analysis this list comprised all 64 linked short variants showing statistical association with MLP formation. During our analysis of CKM subunits, the query list comprised all known short variants (as identified in (42)) from the four CKM genes (*srb10*, *srb11*, *med12*, *med13*). Following the deletion library screen, the input list comprised all known short variants (as identified in (42)) in the 31 hit genes.

### CRISPR-Cas9 gene-editing

Seamless CRISPR-Cas9 gene editing was done using a published protocol (66). Briefly, single-guide RNAs were inserted in the pMZ379 plasmid using a PCR-based method, while homology templates were generated as large primer dimers also using PCR. To design the single-guide RNA and homology template we used the CRISPR4P tool (http://bahlerweb.cs.ucl.ac.uk/cgi-bin/crispr4p/webapp.py). Furthermore, we checked our sgRNAs in Benchling (67) and chose the sequences with the most favourable on-target and off-target scores. The homology templates contained homologous regions at the edges of the gene of interest, allowing for knockouts.

### Gene set enrichment analysis (GSEA)

GSEA was performed with the Bähler lab tool AnGeLi at http://bahlerweb.cs.ucl.ac.uk/cgi-bin/GLA/GLA_input (68). Gene sets included all categories and the significance threshold 0.01 was chosen with FDR correction for multiple-testing.

### RT-qPCR

Cells were grown to exponential phase (OD_600_ ~0.5) and 15ml aliquots were immediately spun down and stored at -80°C. We extracted RNA using a standard hot-phenol protocol (69). We used Turbo DNase (Invitrogen) to digest the residual DNA and performed reverse transcription with the Superscript III kit and oligoDT primers (Invitrogen) according to the manufacturer’s guidelines. We performed qPCR using Fast SYBR Green Master Mix (Applied Biosystems) on a QuantStudio 6 Flex instrument (Applied Biosystems) in fast cycling mode according to the manufacturer’s instructions. Quantification of transcript abundance was done using a relative standard curve. For this, we pooled cDNA from samples expected to have the highest *mbx2* concentration, and created a dilution series with 2x, 2/10x, 2/100x and 2/1000x concentrations. We manually removed values for the 2x concentration, which showed strange amplification patterns for both *mbx2* and *act1* and would have led to an order of magnitude higher inferred *mbx2* fold change values. For the standard curve and each of the three biological replicates, we measured 3 technical replicates.

## Results

### Regulators of MLP formation, but not cell-adhesion proteins, are conserved between fission and budding yeasts

To explore conservation in MLP formation between yeast clades, we identified all genes whose orthologs are annotated to an MLP-related Gene Ontology (GO) term in at least one of *C. albicans*, *S. cerevisiae* and *S. pombe* (Methods). We found that 338 of the annotated gene families (orthogroups) are conserved in all three species (Supp Fig 2A). Intriguingly, however, only one orthogroup was functionally conserved, meaning that the ortholog was annotated to an MLP-related GO term in all three species (Supp Fig 2B). This orthogroup includes the *S. pombe* transcription factor Mbx2 and its orthologs, known as Rlm1p in *C. albicans* and the paralogs, Rlm1p and Smp1p in *S. cerevisiae*. Because the criteria for phenotype annotations are looser than for GO terms, including these annotations might uncover more conserved MLP-related genes. We therefore incorporated annotations from the Fission Yeast Phenotype Ontology (FYPO, (53)) and phenotypic data from *S. cerevisiae* and *C. albicans* (see Methods). Interestingly, only 73 genes were annotated to relevant GO or phenotype terms in *S. pombe*, highlighting the stark knowledge gap in this area compared to *C. albicans* (1035 proteins) or *S. cerevisiae* (1373 proteins). As for GO-term annotations, in this wider set of MLP-related genes, most orthogroups related to MLPs in at least one of the three species were conserved (1259/2096) (Fig 1A, Supplementary Table 5), while only a small subset (18/1259) of those conserved genes were functionally conserved as reflected by MLP-related annotations in all three species (Fig 1B). Besides Mbx2, these included additional proteins involved in transcriptional regulation, parts of the Cdk8 kinase module of the Mediator, and members of the cAMP pathway (Fig 1C).

**Figure 1:**
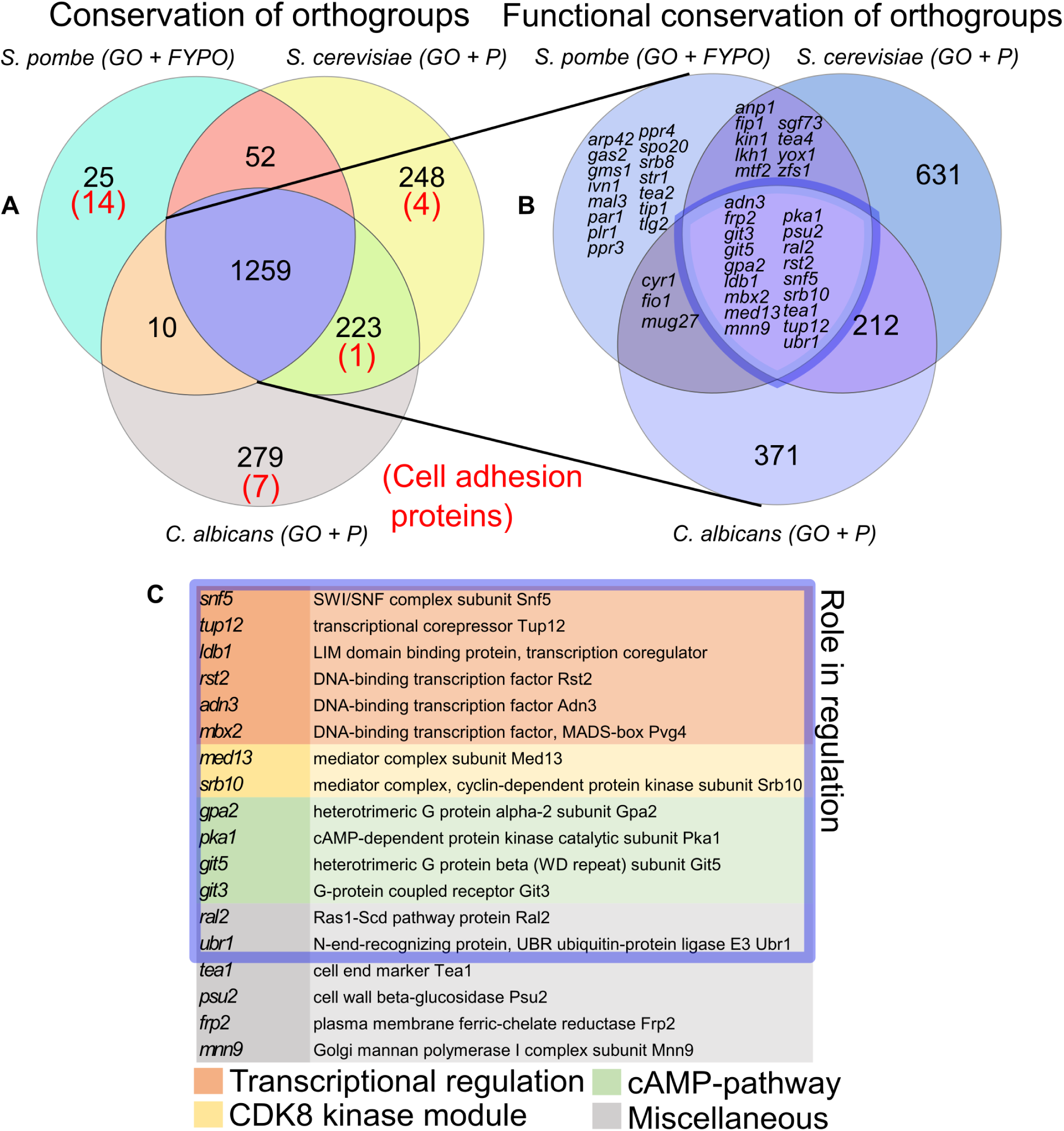
Several regulators of MLP formation are conserved between fission and budding yeast, but cell-adhesion effector proteins are not. A: Venn diagram of numbers of orthogroups, in which at least one gene in the orthogroup is annotated in GO-terms or phenotypic data related to MLP formation in either *S. pombe*, *S. cerevisiae* or *C. albicans*. Red numbers indicate cell-adhesion proteins. B: Venn diagram of orthogroups that are conserved across the 3 species (i.e. the middle subset on A) asking whether they are also functionally conserved, i.e. contain at least one gene that is annotated in GO-terms or phenotypic data related to MLP formation in all three species. C: Functionally conserved genes coloured by their broad functional category as indicated. GO: Gene Ontology, FYPO: Fission Yeast Phenotype Ontology, P: Phenotype annotations. Venn diagrams were made using matplotlib-venn in Python (118).

Partial changes in biological pathways between related species, or biological circuit rewiring, have been well documented (70,71). In most cases this rewiring has occurred at the regulatory level (70), however, our above analysis suggests that in the case of MLP formation, this rewiring appears to have happened at the level of downstream effectors, such as cell-adhesion proteins. Additional support for regulatory conservation comes from work demonstrating that overexpression of the *S. pombe* TF Mbx2 can trigger MLP formation in *S. cerevisiae* (22). Furthermore, overexpression of the *S. cerevisiae* TF Flo8p (*S. pombe* orthologs Adn2 and Adn3) can also trigger MLP formation in *S. pombe* (24). Indeed, *S. pombe* cell-adhesion proteins have little in common with other fungal adhesins (72). *S. pombe* flocculins consist of repetitive beta-sheets and commonly a GLEYA or DIPSY sugar-binding domain at the C-terminus (unlike in other fungal adhesins where similar domains are N-terminal (72)), although the dominant flocculin Gsf2 contains neither of these domains. Gsf2 also seems to be unique to *S. pombe* even within the Taphromycotina lineage, with no detected *S. japonicus* ortholog (50,73). The GLEYA domain is similar to the lectin-like ligand-binding domain of certain *S. cerevisiae* flocculins (72), while the DIPSY-domain has only been identified in species of the Taphromycotina lineage. To verify our orthology analysis, we performed sequence- and structure-based queries using BLAST-P (55) and Foldseek (56), respectively, for *S. pombe* cell-adhesion proteins against all proteins in *S. cerevisiae* or *C. albicans* (Methods). Both sequence- (*C. albicans*, P=1.8E-5; *S. cerevisiae*, P=5.2E-7) and structure-based alignment scores (*C. albicans*, P=0.02; *S. cerevisiae*, P=1.3E-5) were significantly lower for *S. pombe* flocculins compared to queries from a random set of 50 conserved proteins (Supp Fig 2C, Supplementary Table 2), further supporting our observation that cell-adhesion proteins in *S. pombe* are either lineage-specific or weakly conserved.

Taken together, our bioinformatic analysis suggests that, aside from a few key regulatory proteins, most genes involved in MLP formation are not functionally conserved between fission yeast and budding yeast. Some of this discrepancy may be attributed to annotation bias due to less work done in *S. pombe* in this subject. However, the presence of genes that are annotated as contributing to MLP-formation in *S. pombe*, but not annotated as such in budding yeast argues for true divergence. Additionally, cell-adhesion proteins seem to differ greatly between fission yeast and budding yeast. Lastly, the much smaller number of genes annotated to MLP formation in *S. pombe* highlights that these phenotypes are understudied in fission yeast.

### Multicellular-like phenotypes depend on environmental context

We observed that the *S. pombe* natural isolate JB759 (Y0036) sticks to the side of glass flasks and forms clumps in minimal media (EMM) but not in rich media (YES) (Fig 2A). To explore the natural variation in MLP formation across strains and conditions, we developed high-throughput methods to assay flocculation and adhesion to agar (Methods) and applied them to a collection of 57 genetically diverse natural isolates (Fig 2B and 2C, Supp Fig 3A) (42). The two phenotypes were strongly positively correlated with each other (r=0.8, P=4E-14, Supp Fig 3A). This result points to a shared mechanism underlying the two distinct MLPs.

**Figure 2:**
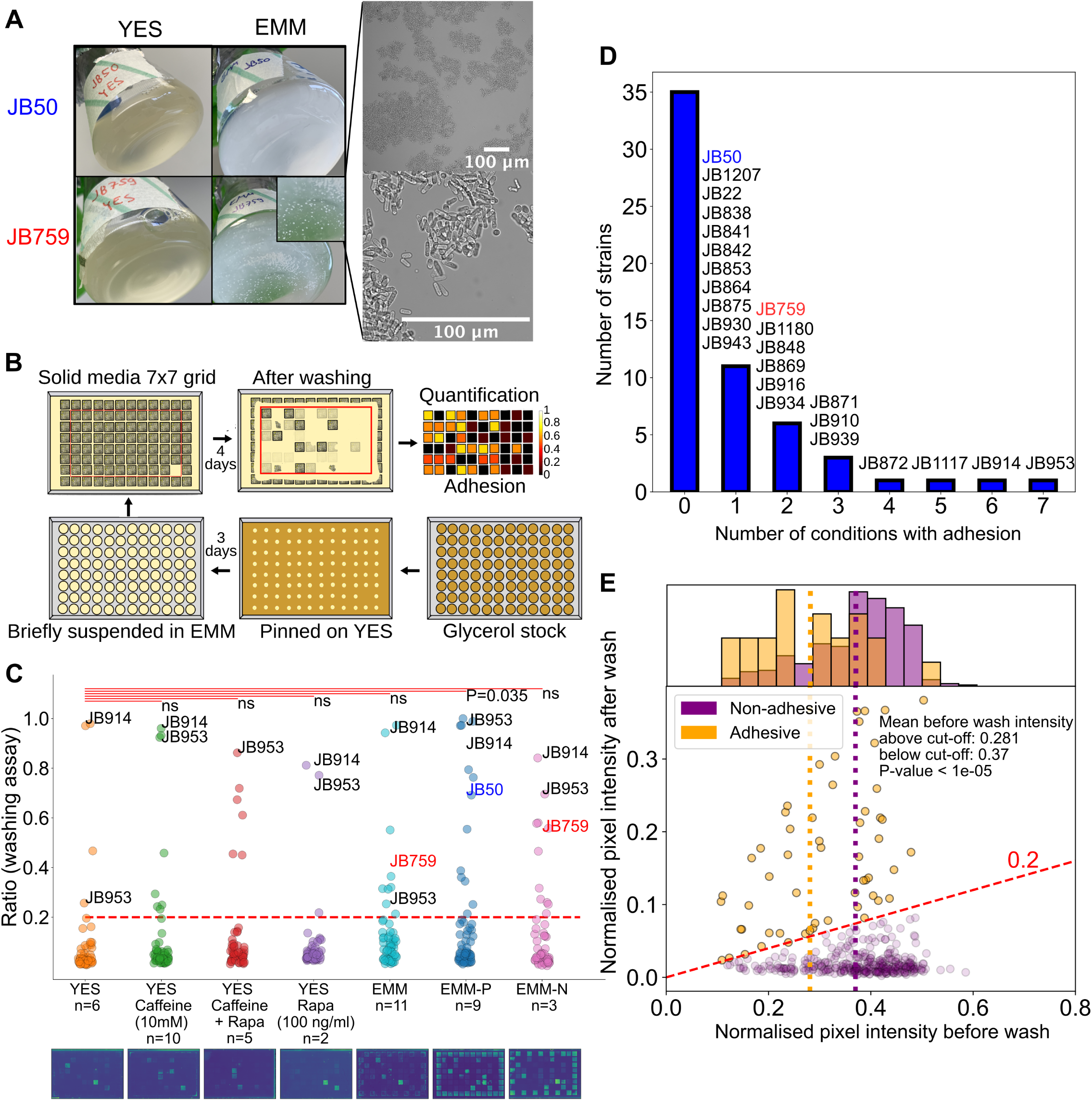
MLP formation in *S. pombe* natural isolates varies with nutrient conditions and is associated with decreased growth. A: Left: Images of our initial observations on the standard laboratory strain JB50 and the natural isolate JB759 showing MLP formation of JB759 in EMM. Right: Microscopy images at two magnifications of the JB759 strain grown in EMM for 2 days. B: Scheme of the high-throughput adhesion assay used to assess MLP formation in *S. pombe.* C: Strip-plot of adhesion to agar across different conditions in the natural isolate library, with images of representative post-wash agar plates below. Each dot represents the mean adhesion value for a given strain in a specific condition. The red dashed line shows a cut-off for strong phenotypes (intensity after wash >0.2 times than before wash). Each condition was compared to rich media (YES) with the null hypothesis that they do not increase MLP formation. P-values were obtained using a one-sided permutation-based T-test and Bonferroni correction. Comparisons were marked not significant (ns.) where the null hypothesis could not be rejected at significance threshold 0.05. Strains around the edges were not taken into account for any statistical analysis (see Methods). The lab strain JB50 and natural isolate JB759 used in panel A are highlighted with colour. D: Histogram showing the number of unique strains forming MLPs in a given number of conditions. E: Scatterplot of mean cell densities before and after washing. Each dot represents one strain in one condition, orange dots represent adhesive data points (ratio of before wash to after wash intensity >0.2, dashed red line) and purple dots represent non-adhesive data points. The histogram shows the distribution of cell densities before washing, as a proxy for growth. The vertical dotted orange and purple lines mark the mean pre-wash densities for the two populations.

Extending our initial observation, depending on the strain, the penetration of surface-adhesion phenotypes varied across different nutrient and drug conditions (Fig 2C, Supplementary Table 6). Compared to adhesion to YES plates, only phosphate starvation (EMM-P) led to significantly increased mean adhesion levels (P=0.035, one-tailed permutation-based T-test). Although the other 6 conditions did not lead to significantly changed mean adhesion levels in the strain collection, between 3 and 13 strains in each condition passed our threshold for a strong adhesion phenotype (defined using an elbow plot, Supp Fig 3B). Compared to the 4 strains that passed that threshold in YES, there were more adhesive strains in EMM-P (n=13), nitrogen starvation (EMM-N) (n=11), EMM (n=9), and YES with caffeine (n=8). Out of the 57 natural isolates, 24 strains (42%) showed a strong adhesion phenotype in at least 1 condition. For most strains, such strong adhesion was limited to 1 or 2 conditions, but 2 strains, JB914 and JB953, showed strong adhesion in 6 or all 7 conditions tested (Fig 2D). Strikingly, even the 2 lab strains, JB22 and JB50, showed strong adhesion under phosphate starvation, while the adhesion level of JB759 was below the threshold in that condition (Fig 2C). Additionally, JB50 exhibited flocculation when grown in EMM-P (Supp Fig 3C). Indeed, a recent RNA-seq dataset from *S. pombe* lab strains grown under similar phosphate starvation conditions reveals that expression levels of the flocculation-related transcription factor gene *mbx2* and downstream cell-adhesion genes increase with time under phosphate starvation (74) (Supp Fig 3D). Though not mentioned in that article, the authors confirmed that “cells started to clump together” under those conditions (Garg, Schwer, Shuman, personal communication). Lastly, comparing measurements across all conditions, we found that strains exhibiting strong adhesion generally featured lower colony density before washing (measured as decreased inverse pixel intensity), which may reflect a decreased growth rate (Permutation-based T-test: P<1E-5, Fig 2E, Supp Fig 3E).

### Truncation of *srb11* causes MLP formation in adhesive natural isolate

Motivated by the findings above, we dissected the genetic mechanism underpinning MLP formation on minimal media in the JB759 strain, originally isolated from wine in South Africa (75). We used an existing segregant library generated from the JB759 strain and the lab strain, JB50 (46) (Fig 3A). Flocculation and adhesion to agar on EMM were highly correlated with each other in this library (Supp Fig 4A, P=2E-20). Notably, adhesion occurred only on EMM and not on YES medium (Permutation-based T-test, P<1E-6; Fig 3B).

**Figure 3:**
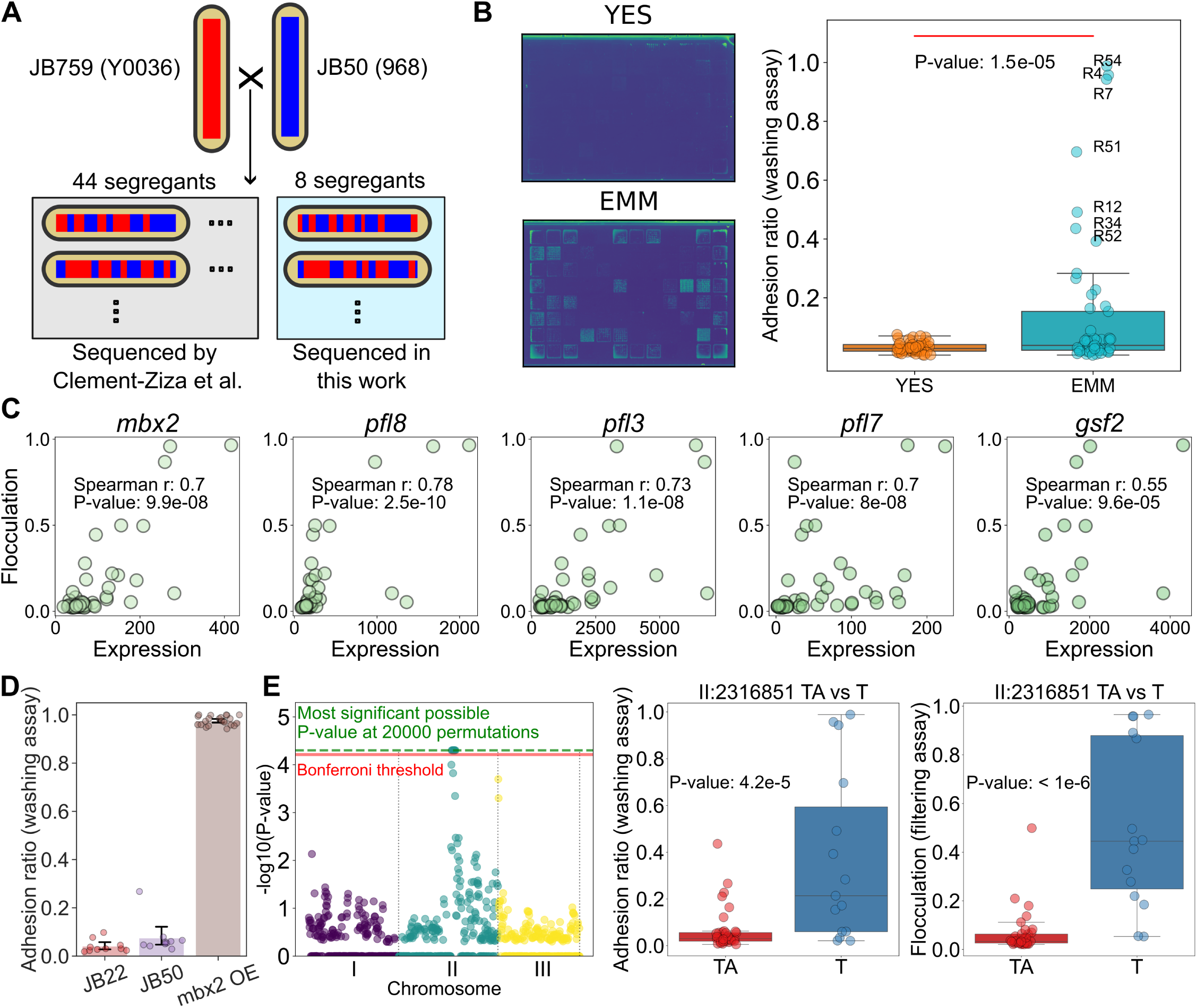
MLP formation in JB759 is driven by *mbx2* expression and associated with a single-nucleotide deletion on chromosome II. A: Scheme for the segregant library. Red and blue stripes indicate genomic recombination resulting from meiosis. SNPs for the strains inside the grey box were previously identified (46) and genome sequencing data for the strains in the light blue box were generated in this work (Methods). B: (Left) Example plates of segregants grown on EMM or YES after washing, shown in viridis colormap. (Right) Adhesion to agar of segregant strains on EMM (mean of 10 replicates) compared to YES (mean of 2 replicates), along with significance of the difference obtained using permutation-based T-test. C: Correlations of *mbx2* and flocculin gene expression with flocculation in EMM. Each dot represents a strain from the segregant library. D: Barplot with measurements overlaid comparing adhesion measurements from standard laboratory strains JB22 and JB50, and the *mbx2* overexpression strain generated in this work (Methods). Error bars represent the 95% confidence interval. E: (Left) Manhattan plot of QTL analysis results for flocculation in EMM. The red line shows the Bonferroni threshold, while the green dashed line shows the highest possible significance achievable using 20,000 permutations. (Middle, Right) Candidate variant is associated with both increased adhesion to agar (mean of 10 replicates) and increased flocculation in EMM (filtering assay, mean of 3 replicates). P-values were determined using a permutation-based T-test with 1E+6 permutations.

An RNA-seq dataset for this segregant library was previously published (56). We used that dataset to perform an unbiased search for correlations between gene expression and flocculation, as measured using our filtering assay (Methods). Following FDR correction, we found 242 genes, of which 138 were protein-coding, to be significantly associated with this phenotype (Supplementary Table 7). The four transcripts showing the highest correlation with flocculation encoded the transcription factor Mbx2 and the flocculins Pfl8, Pfl3 and Pfl7, while the transcript encoding the dominant flocculin Gsf2 was also highly correlated (Fig 3C). Accordingly, *mbx2* gene expression showed a strong association with flocculin gene expression (Supp Fig 4B).

Overexpression of *mbx2* is sufficient to cause flocculation in minimal media (25). We tested whether *mbx2* overexpression also causes agar adhesion detectable by our high-throughput assay. To this end, we engineered the pJR1-41XL plasmid, which contains a *leu* marker and a thiamine-repressible *nmt1* promoter, to overexpress *mbx2* in the leucine auxotroph JB21 strain. Indeed, this *mbx*2 overexpression strain showed strong surface adhesion and flocculation on EMM (Fig 3D, Supp Fig 4C).

To identify genetic determinants of the variation in flocculation amongst the segregants, we mapped Quantitative Trait Loci (QTL) using the available genotype data from 44 of the 52 segregants in the library (56). Our analysis did not allow us to obtain high-fidelity results, with the strongest hit being two SNPs in the non-coding RNA *SPNCRNA.1524*. These variants were also the result of Clement-Ziza et al. (46) for the expression quantitative trait locus of *mbx2*. To increase the statistical robustness of our dataset, we sequenced the remaining 8 strains from the segregant library for which SNPs were not previously identified (Methods). This enabled us to pinpoint a genomic region of 65 short variants that was strongly associated with both flocculation and adhesion to agar (Fig 3E). While these variants did not contain *SPNCRNA.1524*, they did include 26 open reading frames with 13 variants resulting in a changed codon. Among the latter, 1 variant introduced a premature stop codon through a frameshift in the gene *srb11*, encoding cyclin C, and is thus predicted to cause the production of a truncated protein (Fig 4A,B).

**Figure 4:**
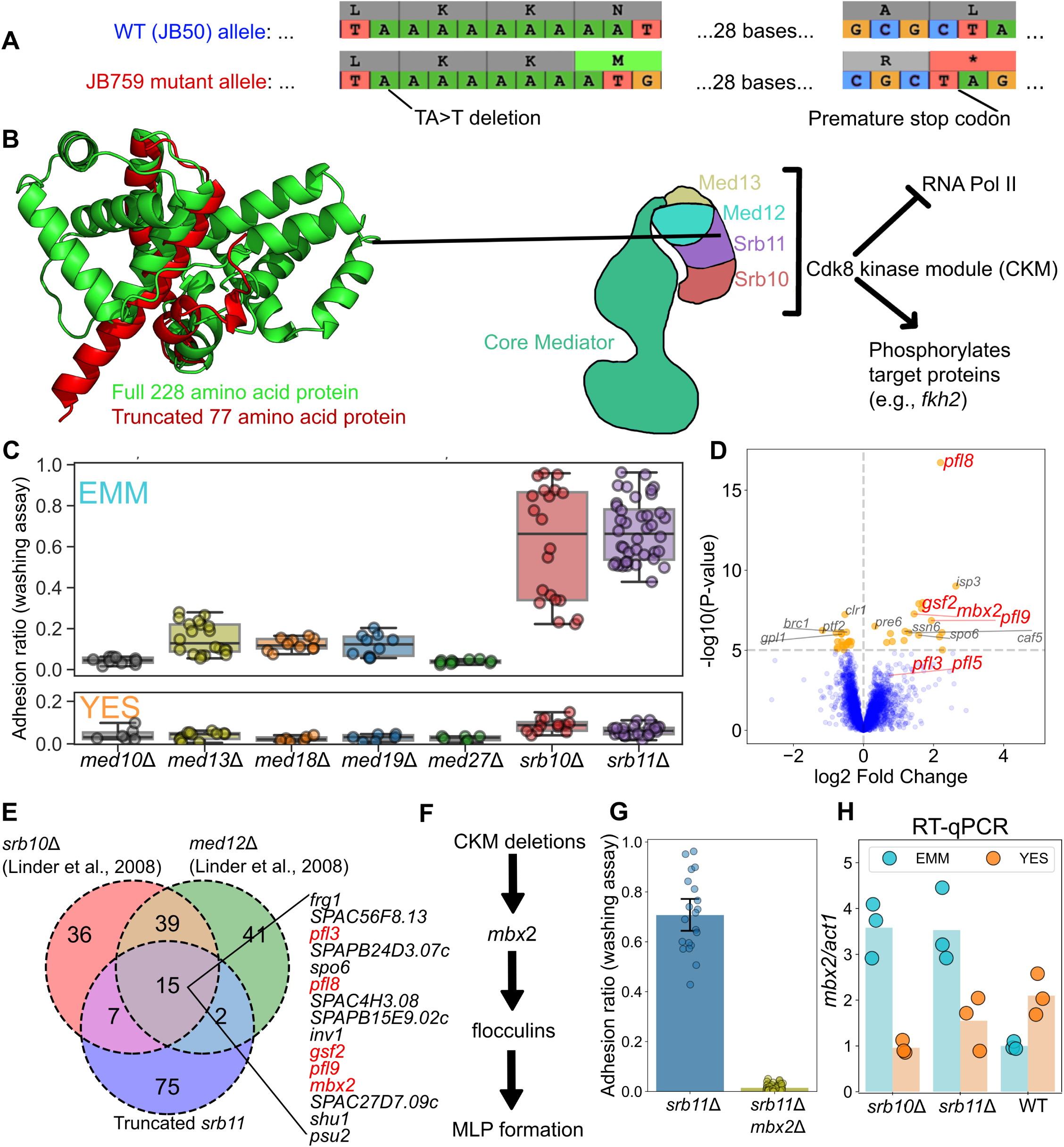
Cdk8 kinase module deletions upregulate *mbx2* in EMM, but not in YES, and lead to MLP formation. A: Scheme showing how a single nucleotide deletion leads to frameshift and premature stop-codon in *srb11*. B: Full Srb11 structure (green) compared with the truncated Srb11 structure (red) as predicted using Colabfold (119). Scheme on the right shows Srb11 in the context of the Cdk8 kinase module of the Mediator complex. The structure was sketched based on structural data (77), and functional roles were summarized based on (77,79). C: Strip-boxplot showing adhesion values from Mediator gene deletion strains on EMM and YES as indicated. Each dot represents a replicate. D: Differential expression analysis of segregant strains split on the II:2316851 TA>A single-nucleotide deletion. Fold-change values and P-values were obtained from DESeq2 (59). E: Overlap of upregulated genes in three CKM mutants based on our *srb11* data and data from Linder et al. (60). F: Simplified model for how CKM deletions lead to MLP formation. See main text for details. G: Adhesion measurements for the *srb11Δ* strain obtained from the deletion collection, and its derived strain after *mbx2* knock-out with CRISPR. Each dot represents a replicate measurement. Error bars represent the 95% confidence interval. H: RT-qPCR showing *mbx2* expression of *srb10Δ::Kan*, *srb11Δ::Kan* and wild-type (WT) strains in EMM or YES. Height of each bar reflects the mean of three biological replicates which are indicated by dots.

Cyclin C (Srb11), together with cyclin-dependent kinase 8 (Srb10) and Mediator subunits Med12/Srb8 and Med13/Srb9, is part of the Cdk8 kinase module (CKM) of the Mediator complex (Fig 4B). The CKM can impede transcription by inhibiting the interaction between the core Mediator and RNA polymerase II (76–78), and phosphorylates various target proteins (79–83). Previously it was found that deletion of *srb10*, *med12* and *med13* causes flocculation (60,84), which was supported by microarray data from Linder et al. (60) showing increased expression of flocculins in these mutants.

To better understand the role of the Mediator complex in cell adhesion, we studied deletion mutants of all subunits of Mediator available in the prototrophic gene-deletion library (43,44) (Supplementary Table 8). In accordance with previous work (60,85), we found that the *srb10*Δ strain flocculated (Supp Fig 5A) and exhibited adhesion to agar, particularly on EMM (Fig 4C). Additionally, *srb11*Δ also exhibited strong flocculation and surface adhesion phenotypes on EMM. The *med13Δ* strain showed a milder phenotype compared to the extreme adhesion of *srb10Δ* and *srb11Δ* cells (Fig 4C). The *med12* gene was not represented in the deletion library but, similar to *med13Δ,* it was previously noted that *med12Δ* cells show a milder flocculation phenotype compared to *srb10Δ* cells (60). Additionally, deletion of the core Mediator genes *med19/rox3* and *med18* also resulted in mild adhesion phenotypes, while deletion of *med10/nut2* and *med27/pmc3* did not (Fig 4C). We then checked whether flocculation or adhesion to agar in the CKM mutant strains depends on media composition. Interestingly, the core Mediator mutants only showed their adhesion phenotype on EMM, while the CKM mutants (*srb10Δ*, *srb11Δ* and *med13Δ*) showed a strong phenotype in EMM as well as a very mild adhesion phenotype on YES (Fig 4C).

To further validate whether the lack of *srb11* is sufficient to cause MLP formation in our non-adhesive parental strain (JB50), we independently knocked out *srb11* using seamless CRISPR-Cas9 gene-editing (66) (Supp Fig 5B). The resulting *srb11Δ* strain was slightly more adhesive than the *srb11Δ*::*Kan* strain from the prototrophic deletion library (Supp Fig 5C), likely because the deletion library strain has accumulated suppression mutations in other genes.

Next, to identify upregulated genes in strains containing the *srb11* truncation, we first split the segregants by their *srb11* haplotype, and then performed differential expression analysis using the RNA-seq dataset from Clement-Ziza et al., which was generated from the same segregant library grown in EMM (46) (Fig 4D, Supplementary Table 9). The upregulated genes were compared with those in the *srb10*Δ and *med12*Δ strains examined by Linder et al., which were also generated from strains grown in EMM (60). We found that *mbx2* and four flocculin genes (*pfl3, pfl8, gsf2,* and *pfl9*) were upregulated in all 3 mutants, together with 10 other genes (Fig 4E). The latter included the *inv1* gene for external sucrose invertase whose ortholog in *S. cerevisiae, SUC2*, can enable nutrient sharing among cell aggregates (8,16). Furthermore, the cell surface heme acquisition gene *shu1* causes flocculation and filamentous growth when overexpressed (30).

We then asked how many of the genes upregulated in the CKM deletion mutants overlap with genes known to be activated by Mbx2. Based on microarray and ChIP-chip data from Kwon et al. (25), the four flocculins and *inv1* are indeed regulated by Mbx2 (Fig 4F, Supp Fig 6A). To test whether MLP formation in the *srb11*Δ strain requires *mbx2*, we created an *srb11/mbx2* double mutant using seamless CRISPR-Cas9 gene-editing (66) (Supp Fig 5B). This double-mutant strain exhibited no flocculation in liquid media (Supp Fig 6B) or adhesion to agar (Fig 4G). We conclude that Mbx2 is essential for the MLP phenotype seen in *srb11*Δ cells.

Given the model that the *srb11* truncation upregulates *mbx2*, we wondered why the JB759, *srb10*Δ and *srb11*Δ strains only exhibited strong MLP formation in EMM. The transcriptomics data from the segregant library (46) and the CKM deletion strains (60) both came from cells grown in EMM. We, therefore, tested whether the upregulation of *mbx2* in CKM deletion strains is specific to EMM, like MLP formation. To this end, we performed RT-qPCR in wild type (JB50) and CKM deletion strains (*srb10*Δ, *srb11*Δ) from the deletion library grown in EMM or YES to measure the expression of *mbx2*. The upregulation of *mbx2* in the CKM mutants was indeed exclusive to EMM (Fig 4H). The 3.5-fold upregulation of *mbx2* in CKM mutants in EMM was similar to the increase observed using microarray data from an *srb10*Δ deletion strain (9.3-fold, (60)) or when analyzing bulk segregant data averaged over *srb11* truncation haplotypes (Fig 4D, 2.7-fold, (46)).

Although the different proteins of the *S. pombe* CKM physically interact (84,86), and their mutants feature similar phenotypes and transcriptomic profiles in *S. pombe* (72) and *S. cerevisiae* (87), here we show that under our growth conditions, deletion of individual parts of this subunit leads to strikingly different adhesion phenotypes. To see whether this is also true for other phenotypes, we analysed data from Rodriguez-Lopez et al. (88), who measured sensitivity and resistance phenotypes of deletion strains in 131 conditions. Deletion of *med13* and *srb11* results in different phenotypes across a range of growth conditions. In terms of these phenotypes, *med13Δ* is not more similar to *srb11Δ* than expected by chance (Supp Fig 7). Interestingly, amongst the top-10 deletion strains that were phenotypically most similar to *srb11Δ,* we identified *ace2Δ* and *cbf11Δ*, both of which have been found to trigger MLP formation (26,60) (Supp Fig 7).

The only well-documented physical interaction of Srb10/Srb11 in *S. pombe* is the stabilisation of the transcription factor Fkh2 by phosphorylation (79,80). We therefore looked at whether loss of *fkh2*, similarly to loss of *srb10/srb11,* leads to *mbx2* upregulation. ChIP-seq data (89) and our analysis of microarray data (79,89) indicated that Fkh2 does not bind to the *mbx2* promoter, nor does deletion of *fkh2* increase levels of *mbx2* (Supp Fig 8A). The mechanism that inhibits the upregulation of *mbx2* in YES also remains unknown, as its only known repressor, *gsf1*, is not significantly upregulated in rich media relative to minimal media ((90), Supp Fig 8B).

Finally, we checked whether similar nonsense mutations appear in any other CKM genes within the natural isolate library using the genotype data from Jeffares et al. (42). Besides JB759, the parental strain of the segregant library, we found the same *srb11* frameshift mutation to appear in the unrelated, strongly flocculant strain JB914. Interestingly, however, this strain exhibited strong adhesion to agar (and flocculation) on both EMM and YES (Fig 2C,D), indicating the likely presence of one or more mutations in the pathway that inhibits MLP formation in YES.

In summary, we used a segregant library to dissect the genetic determinants of MLP formation on EMM in the JB759 strain. We found a single-nucleotide deletion that leads to a truncation of Srb11 to be associated with MLP formation on EMM, and determined its effect to be the EMM-specific upregulation of the transcription-factor gene *mbx2*. Upregulation of *mbx2* in turn leads to the upregulation of cell-adhesion genes which mediate MLP formation. Using an *mbx2* overexpression strain and an *mbx2*/*srb11* double mutant, we showed that upregulation of *mbx2* is both necessary and sufficient to explain MLP formation in the *srb11* mutant JB759.

### Novel players in MLP formation on minimal media

The premature stop codon in *srb11* only accounts for the phenotype of two strains (JB759 and JB914) out of the 7 natural isolates we have found to exhibit MLP formation on EMM. Therefore, to identify further possible genetic causes of agar adhesion on EMM, we screened the prototrophic gene-deletion library (43,44) and a recently created lincRNA deletion library (45). We performed one round of the adhesion-to-agar assay on EMM for 3721 unique deletion strains (a total of 4327 strains including replicates). While not every strain grew on YES after initial inoculation, or grew on EMM during the screen, we successfully assayed a total of 3628 strains (Fig 5A). The *srb10Δ* and *srb11Δ* strains failed to be inoculated for this initial screen, likely due to aggregation at the bottom of the 96-well plate. This indicates that some other true positive genes related to MLP formation may have been missed in our initial screen.

**Figure 5:**
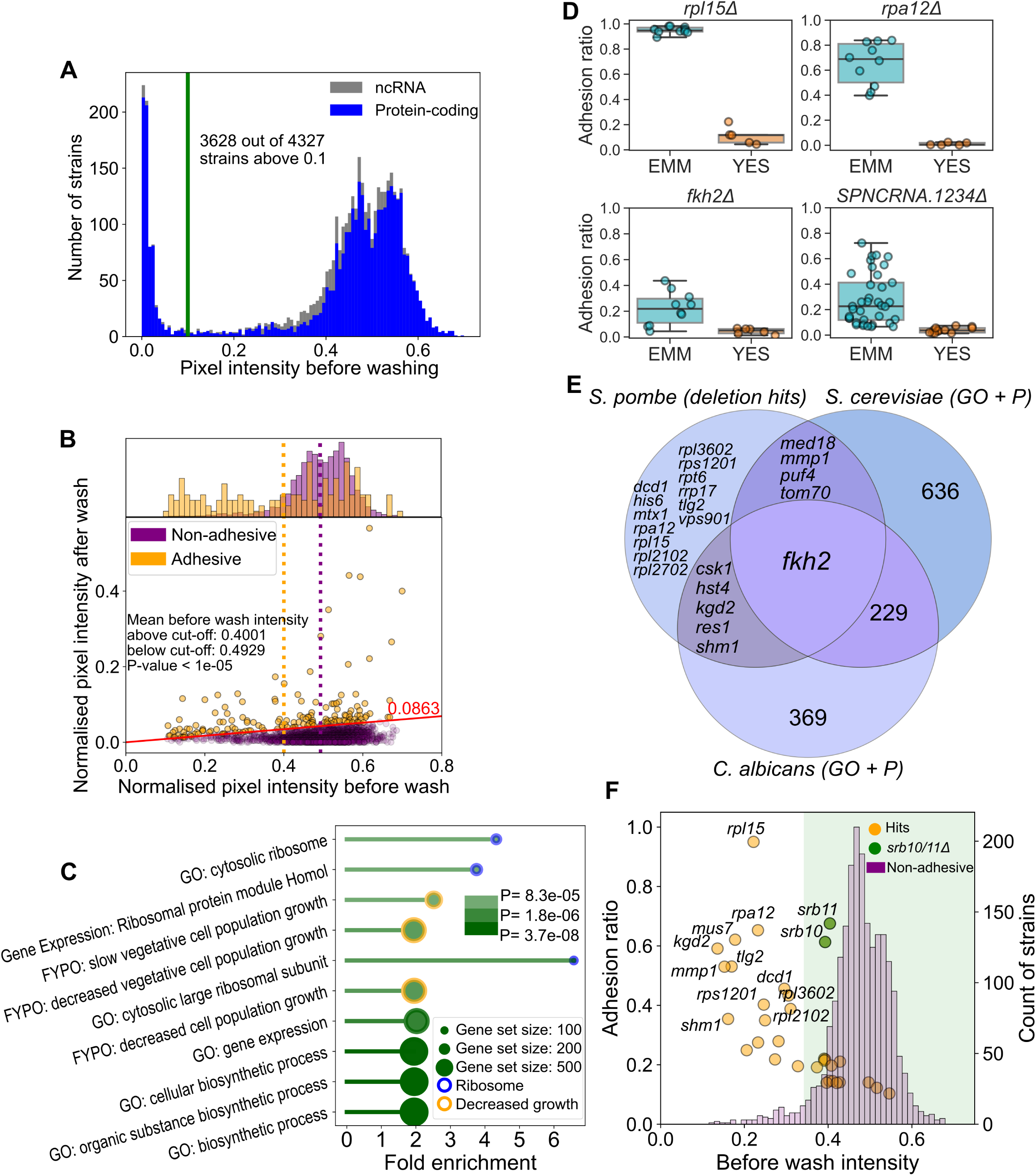
Deletion library screen identified 31 genes associated with MLP formation on EMM. A: Histogram showing cell densities of strains in our deletion library screen before washing. B: Scatterplot of mean cell densities before and after washing. Each dot represents a deletion strain, and colors represent adhesive and non-adhesive strains as indicated. The red line represents the cut-off at the 95th percentile of adhesion values, also shown on Supp Fig 9. The histogram shows the distribution of cell densities before washing as a proxy for growth. The dotted orange and purple lines mark the mean pre-wash intensity values for the two populations. The P-value was determined using a permutation-based T-test with 1E+5 permutations. C: Barplot showing the fold enrichment of the top-10 most significantly enriched processes, with blue circles for terms associated with ribosomes and orange circles for terms associated with decreased growth. Terms were sorted based on P-values and increasing color intensity represents increasing -log10(P-value). The size of the circle at the end of each bar represents the size of the gene set. D: Strip-boxplots of adhesion ratios obtained with the washing assay for 4 of the 31 verified hits on EMM (light blue) vs YES (orange). Each dot is an independent observation. E: Venn diagram showing the functional conservation of the genetically conserved hits from our screen. Only *fkh2* is annotated as being involved in MLP formation in all three species. Venn diagram was made using matplotlib-venn in Python (118). F: Scatterplot of adhesion ratios and before-wash colony intensities overlaid by a histogram showing before-wash colony intensities of non-adhesive deletion strains which were assayed in the middle 60 spots during the original screen. The green shaded area marks strains that are above the 5th percentile of colony intensities in the non-adhesive strains. The *srb10/11*Δ strains, highlighted with green, are the most adhesive strains from those with growth values above the 5th percentile.

Given that our measure for adhesion is the fraction of cells remaining after washing (after/before), we worried that strains with minimal growth before washing (denominator) might appear to have higher ratios, despite only negligible intensity values after washing (attributed to measurement error rather than biological signal). A scatterplot of all of our measurements argues against such systematic bias, as adhesive strains cover a wide range of pre-wash growth values (Fig 5B). Still, the adhesive deletion strains exhibited a decreased growth phenotype on average (Fig 5B, Permutation-based T-test, P<10E-5), but we attribute this to a biological effect similar to that seen in the natural isolates (Fig 2E). Based on the assay, deletion strains of protein-coding genes with adhesion values in the top-5 percentile were chosen for functional enrichment analysis (Supp Fig 9). These mutants showing the strongest adhesion phenotypes were enriched for ribosomal protein genes and for genes associated with slow-growth phenotypes (Fig 5C, Supplementary Table 10, 11).

To validate these findings, we narrowed down our search to the most adhesive strains. By arranging them in the middle 60 spots of three 96-well plates and including a positive control (strongly adhesive JB914 strain) and negative controls (non-adhesive deletion strains and the lab strain JB50, as well as an empty square), we were able to quantify the adhesion of these strains more precisely. In this confirmation step, we found 31 high-confidence hits (Supplementary Table 12), defined as deletion strains where, in at least 5 repeats, cell density before (>0.1 normalised pixel intensity) and after washing (>0.05 normalised pixel intensity) was sufficient to allow robust quantification of adhesion, and the adhesion ratio was greater than 0.086, the 95th percentile cutoff for our initial screen. Interestingly, except for the *sre2* mutant, all adhesion phenotypes were either milder or not present on YES (Fig 5D, Supp Fig 10).

Out of these high-confidence hits, *sre2*, a sterol regulatory element binding transcription factor (25), *rpl2102,* a part of the large ribosomal subunit (38), and *med18*, a component of the Mediator head domain (60,91), have been previously implicated in cell adhesion or filamentous growth. We found three lincRNA deletions to exhibit adhesion, but at least 2 of these likely affect protein-coding genes: *SPNCRNA.1234* entirely overlaps with the gene *nmt1*, while *SPNCRNA.781* is near the promoter of *hsr1*, a transcription factor which was recently identified in our lab to bind promoters of flocculin genes (unpublished ChIP-seq data; Olivia Hillson). The third lincRNA, *SPNCRNA.900*, is placed between two genes, *glt1* and *eme1*, neither of which appears to be obviously related to MLP formation, and therefore could be a *bona fide* trans-acting non-coding RNA that influences MLP formation.

We then asked whether these hits might be belong to the same pathway or represent separate triggers for MLP formation. To answer this, we returned to the large-scale phenotypic dataset from (88) (analysed in the previous section), where the authors identified 8 broad phenotypic clusters of deletion strains. We found our hits to be spread out amongst clusters, as they were present in 6 out of 8 such groups (Supp Fig 11). This suggests that while these gene deletions all lead to MLP formation on EMM, they represent different pathways, and their deletions lead to different phenotypes across conditions.

We looked at whether these hits are also functionally conserved in *C. albicans* and *S. cerevisiae*. Four protein-coding genes (*SPAC607.02c*, *sre2, meu27, for3*) do not have an ortholog in the two budding yeast species. From the orthogroups that are genetically conserved, ribosomal gene deletions only affected MLP formation in *S. pombe*, while some orthogroups contained genes related to MLP formation in *C. albicans* (*csk1*, *hst4*, *kgd2*, *res1*, *shm1*) or *S. cerevisiae* (*med18*, *mmp1*, *puf4*, *tom70*). Only one gene, *fkh2* seems to have a conserved role across all three species in the regulation of MLP formation (92–94) (Fig 5E).

Finally, we checked whether mutations of these hit genes appeared in any of the natural isolates. The number of non-synonymous SNPs normalised by total SNPs in our 31 genes was slightly higher in wild isolates that showed adhesion in EMM vs wild isolates that did not show adhesion (Permutation-based T-test, P=0.02, Supp Fig 11B). These variants and other synonymous mutations, or mutations in regulatory regions could contribute to MLP formation in wild isolates, however none of the 31 genes carried a more severe nonsense or frameshift mutation. Given that such mutations would lead to decreased growth efficiency, it is not surprising that they are absent in natural isolate genomes. This, however, raises the question of what makes the *srb11* null mutation so special that it appears in two unrelated natural isolates (based on phylogeny in (42)). To answer this question, we examined the trade-off in MLP formation versus growth efficiency across all our measurements, revealing that *srb10* and *srb11* deletions present an ideal combination of strong adhesion (2nd and 5th most adhesive) and growth efficiency (both above 5th percentile of all non-adhesive deletion strains) compared to other adhesive deletion strains (Fig 5F).

In summary, our deletion library screen for EMM agar adhesion identified 31 high-confidence hits, including genes unique to *S. pombe* as well as genes that may be functionally conserved in that they are annotated as contributing to MLP formation in budding yeasts. Additionally, we identified the *srb11* null mutant to provide higher adhesion while maintaining better growth efficiency than these hits, possibly explaining its presence as a natural QTL.

## Discussion

### MLP formation as an adaptation to environmental conditions?

Although *S. pombe* has been a popular model organism for decades, its multicellular-like phenotypes (MLPs) have received little attention. We find that many natural isolates exhibit MLP formation, indicating that MLPs play an important role in the natural ecology of the species. While MLPs have been understudied in *S. pombe,* there has been much more work in the budding yeasts *S. cerevisiae,* where flocculation is important for winemaking and brewing (9), and *C. albicans,* where biofilm formation has been linked to pathogenesis (14,15). Comparing the genes associated with MLP formation between these three species revealed several conserved proteins that regulate MLP formation, while effector cell-adhesion proteins are not conserved. The rapid evolution of cell-adhesion proteins has been noted before (72,95–97), suggesting a possible role in adaptation to new environments and divergence of cell-cell interactions, and possibly contributing to speciation. However, it remains unclear to what extent differences in cell-adhesion proteins limit interactions between strains and even species, as different yeast species can co-flocculate (98,99). Regardless, the contrast between highly variable effector proteins and conserved regulatory proteins is striking given that evolution is known to rewire regulatory interactions while maintaining stable effector proteins in other pathways (70).

Although some regulatory proteins are conserved across species, their activity likely varies even within species across different conditions. Concentrations of minerals (e.g., Ca2+ (24)) and pH (30) can also directly affect the function of cell-adhesion proteins. Specifically, in the case of adhesion to agar, we show that different *S. pombe* strains exhibit MLP formation under different nutrient conditions (Fig 2A). While there may not be a single environmental trigger for MLP formation across strains, 42% of the natural isolates showed strong MLP formation on at least one growth media, and phosphate starvation generally triggered the largest changes across all strains. Still, it is not clear what advantage these phenotypes confer to *S. pombe* cells, and why they vary so widely between strains. One possibility is that, similarly to *S. cerevisiae* (19), aggregation in *S. pombe* allows cells to thrive in nutrient-poor environments by increasing local nutrient concentrations through a shared pool of excreted enzymes. Indeed, we found that *mbx2* upregulation, caused by CKM deletions, results in significant upregulation of the invertase gene *inv1*, which might facilitate the sharing of digested monosaccharides as “public goods” (16,19). A similar external enzyme, the acid phosphatase Pho1, participates in phosphate scavenging during phosphate starvation (100), suggesting an experimentally testable selective advantage of MLP formation in that condition.

### Cyclin C: genetic insight into the natural variation in MLP formation

In addition to the effects of environmental conditions, the genetic basis of natural diversity in MLP formation of *S. pombe* was also poorly understood. We find that the South African strain JB759 exhibits moderate levels of MLP formation, and that this phenotype is driven by a truncation of *srb11,* encoding cyclin C, a component of the Cdk8 kinase module (CKM) of the Mediator. The canonical function of the Mediator complex is to form a bridge between general transcription factors and RNA polymerase II (Pol II), and this complex is highly conserved across eukaryotes, from yeast to humans (77). The Mediator has four key subunits, the head, middle and tail modules forming the core Mediator, and the CKM can reversibly bind to this core (60,76–78). Our key finding is that in minimal media, loss of CKM results in upregulation of many genes, one of them encoding the transcription factor Mbx2, which then activates expression of the flocculins as well as other genes, e.g. the external sucrose invertase *inv1* (Fig 4). The canonical function of the CKM is to inhibit Pol II recruitment to the promoter, and thereby to repress basal transcription (76,77). However, such transcriptional repression is not thought to be gene-specific, as the DNA-binding profiles of the CKM match the broad DNA-binding profiles of the core Mediator (78). Several instances of non-canonical functions of the CKM have been found in yeast species. In *C. albicans,* the CKM phosphorylates the hyphal growth promoting transcription factor Flo8p, which is thereby targeted for degradation, thus repressing hyphal growth and adhesion (81). In *S. cerevisiae,* the CKM can affect histone lysine methylation and repress the expression of the cell-surface flocculin gene *FLO11* and of the *inv1* ortholog, the sucrose invertase gene *SUC2* (101). Furthermore, in *S. cerevisiae,* the CKM phosphorylates the transcription factors Ste12p and Phd1p (82,83), leading to their degradation and repression of filamentous growth. The CKM therefore seems to be a conserved repressor of MLP formation across yeast species.

In *S. pombe,* the most studied aspect of the CKM is its regulation of mitotic entry through periodic phosphorylation of the forkhead transcription factor Fkh2 (79). Interestingly, *fkh2* came up as a hit in our deletion screen, and its adhesion phenotype was also EMM-dependent, although milder than that of *srb10* and *srb11* deletions. Orthologs of *fkh2* are negative regulators of MLP formation in *C. albicans* and *S. cerevisiae* as well (92–94). In *S. pombe*, the phosphorylation of Fkh2 by Srb10 inhibits its degradation, and therefore allows Fhk2 to accumulate and trigger entry into mitosis (79). Surprisingly, while a lack of Srb10 activity delays mitotic entry, deletions of *med12/srb8* or *med13/srb9* show the opposite phenotype, advancing mitotic entry (80). The authors’ explanation is that normally Med12 and Med13 anchor Srb10 and Srb11 to the Mediator, and deletion of this anchor results in an active pool of free Srb10 and Srb11, ready to phosphorylate Fkh2 (80). MLP formation seems to be another such phenotype that differs strikingly between deletions of different parts of the CKM, as *med13Δ* (and *med12Δ* (60)) leads to milder adhesion phenotypes compared to *srb10Δ* or *srb11Δ*. Surprisingly, *mbx2* transcript levels seem to be upregulated in all CKM deletions for which transcriptomic data exists (*srb10Δ, srb11Δ,* and *med12Δ,* see Fig 4E), and it is unclear what makes their phenotypes different. It also remains unknown how exactly the CKM affects Mbx2. Given the non-canonical roles of the CKM in *S. cerevisiae* and *C. albicans* (81–83), a possible scenario is that it phosphorylates Mbx2, which results in its degradation (Fig 6). In this case, deletion of the CKM would result in the accumulation of Mbx2, which binds its own promoter (25) and would therefore trigger upregulation of the *mbx2* transcript. Alternatively, the CKM might phosphorylate and stabilise a repressor of *mbx2*. Further dissection of this pathway will require phosphoproteomic data similar to that recently collected in *C. albicans* (81). It also remains unclear why CKM deletions result in the upregulation of *mbx2* only in minimal medium, suggesting a repressive mechanism in rich medium (Fig 6). Further investigation of the natural isolate JB914 which contains the *srb11* truncation while showing MLP formation in both rich and minimal media might help identify such a mechanism.

**Figure 6:**
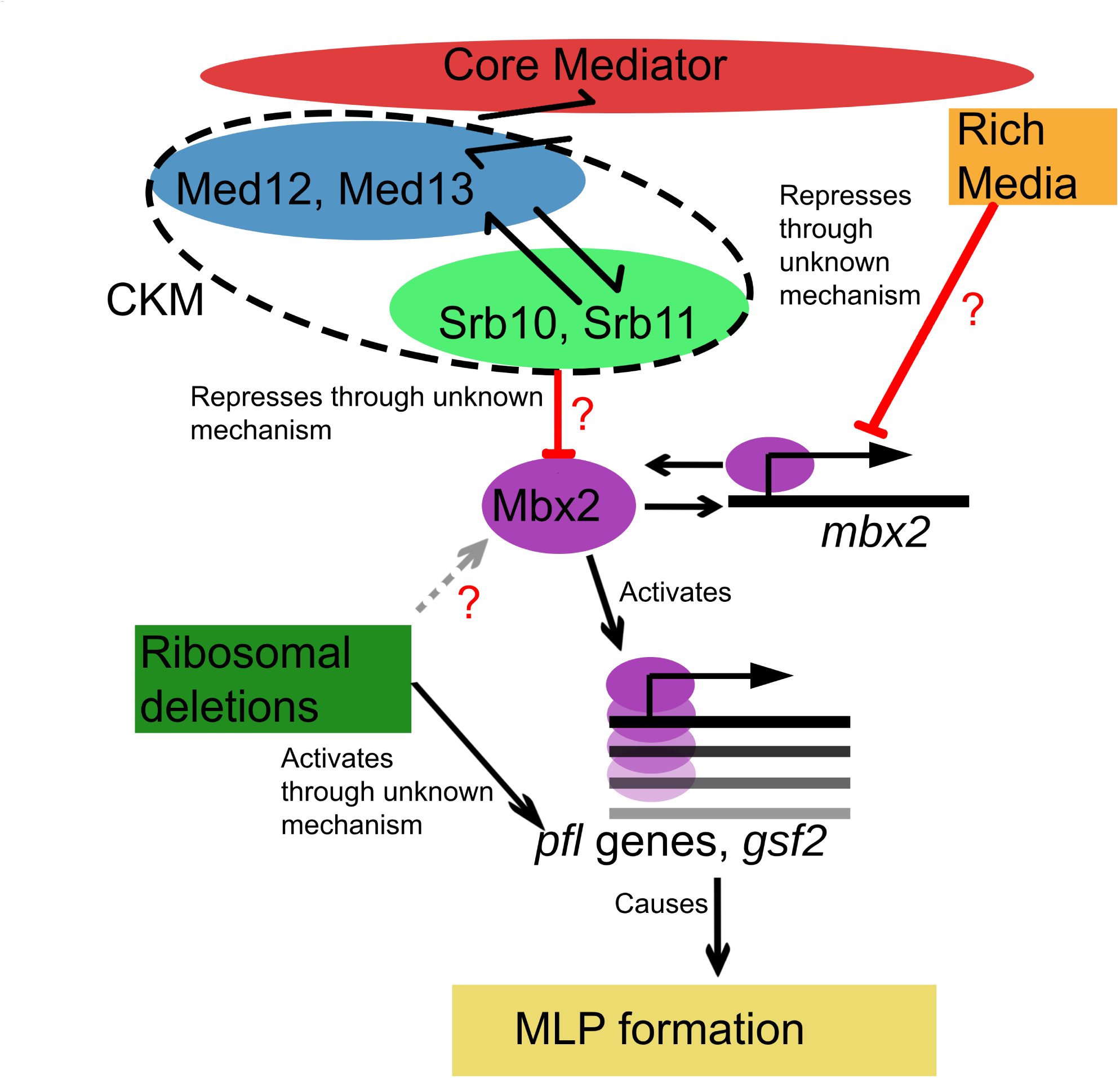
Model for EMM-dependent MLP formation of CKM mutants. We propose that the CKM phosphorylates Mbx2 and targets it for degradation, based on similar observations in *C. albicans* (81) and *S. cerevisiae* (83). Additionally, the *mbx2* transcript is repressed through an unknown mechanism in YES as we observed using RT-qPCR. In minimal media, if members of the CKM are deleted, *mbx2* becomes upregulated, triggering the expression of flocculin genes, which in turn cause MLP formation. Deletion of ribosomal genes also triggers MLP formation, although it is unclear whether this occurs through upregulation of *mbx2* or directly through the flocculin genes. The coloured boxes show our main pathway of interest. The red arrows show our main findings, while red question marks show the main outstanding mechanistic questions.

Besides the extreme adhesion phenotypes of CKM mutants, we find that deletion of genes for two other mediator complex subunits, *med18* and *med19/rox3,* also cause mild adhesion to the agar surface. A positive genetic interaction has previously been found between *mbx2* and *med19/rox3*, supporting our finding (102). In addition to adhesion phenotypes, Mediator head mutants (including *med8, med17, med18, med20,* and *med27* deletions) display filamentous growth, an MLP that reflects a lack of cell separation after mitosis. This phenotype occurs through loss of expression of the transcription factor *ace2* (60). Interestingly, cell separation is also regulated by Ace2 in *S. cerevisiae*, and various Mediator defects cause a drop in transcription of Ace2 targets in that *S. cerevisiae* (60). From all these findings, the Mediator emerges across divergent yeast lineages as a conserved central hub of MLP regulation, upstream of Mbx2 which drives expression of cell-adhesion proteins that cause separated cells to adhere to one another, and/or Ace2, which prevents cells from separating following division, thus driving filamentation.

Although we identified 31 genes whose deletion results in adhesion, as for *srb11*, none of the natural isolate strains carried a null mutation in those genes, unlike for *srb11.* We hypothesize that the benefit of *srb10* or *srb11* deletions, compared to deletions in the other genes, lies in their strong adhesion phenotypes coupled with only a slight compromise in cell growth. Therefore, if MLP formation in a low-nutrient environment is selected for, a null mutation in *srb10* or *srb11* might be one of the most favourable outcomes of sampling genotypic space by random mutations.

### Ribosomal genes and MLP formation: A novel pathway?

The 31 hits from our screen include 5 ribosomal genes (*rpl15, rpl3602, rps1201, rpl2102, rpl2702*) (Fig 6). Each of these genes has a paralog, as most ribosomal genes in *S. pombe* do, and is therefore likely somewhat dispensable. Ribosomal paralogs are tuned to certain translational responses in *S. cerevisiae* (103–105). In *S. pombe*, there is also evidence for paralog-dependent differences in ribosome compositions (106). MLP formation caused by the deletion of these genes might be unique to *S. pombe*, as this phenomenon has not been reported in *S. cerevisiae* or *C. albicans*. This idea also fits the observations of Li et al. (38) who found that deletion of *rpl3201*, *rpl3202*, or *rpl902* also cause MLP formation in *S. pombe* (together with our high-confidence hits a total of 8 ribosomal genes). Liu et al. (39) then linked the *rpl3201* and *rpl3202* deletions to the upregulation of the flocculin genes (Fig 6), which might be mediated by Mbx2 in this case as well. In our screen, *rpl3202Δ* exhibited adhesion above the 95th percentile, however its post-wash intensity was below our threshold for robust quantification in the verification step, likely due to impaired growth, and it was therefore filtered out from our final hits (similar to two additional ribosomal deletions: *rpl2101Δ* and *rpl3702Δ*). Nitrogen starvation and addition of caffeine are strongly linked to decreased translation (49), and we identify several natural isolates in which those conditions trigger MLP formation (Fig 2C). They may cause MLP formation through a similar pathway to that triggered by the deletion of these MLP-related ribosomal genes. Such ribosomal deletions might mimic physiological circumstances of low levels of translation (e.g., inhibition of ribosomes due to toxins or starvation). Under such circumstances, individual cells might not be able to produce sufficient amounts of specific proteins that repress MLP formation. Alternatively, the missing ribosomal subunits could lead to metabolic triggers that cause the cell to sense starvation. In the latter case, forming MLPs might be an adaptive strategy that allows starving cells to share “public goods”, e.g., extracellular enzymes and metabolites. If this is a general mechanism that results from proteome-wide decreases in translation, it is unclear why only certain ribosomal subunits triggered MLP formation in our screen, when a total of 98 ribosomal gene deletions (as captured by GO:0005840 ribosome) were assayed. Synthetic genetic arrays (107) using these ribosomal deletions as query strains, and assaying adhesion to agar on minimal medium, could uncover potential members of this new pathway linking translation levels to MLP formation.

### Final remarks

It is often implicitly assumed that yeast colonies are a homogeneous mass of unicellular organisms. It has recently become clear, however, that there is considerable heterogeneity between genetically identical cells (108,109). For example, single-cell RNA-seq data indicate that *S. pombe* cells under limiting glucose feature highly variable gene expression across cells as growth decreases (109). When cells form MLPs, such heterogeneity might be amplified by differential access to nutrients or exposure to stress based on a cell’s position within the floc or filament. Our understanding of stress or starvation responses may underestimate the role of MLPs and the phenotypic heterogeneity that they generate. First, most work on stress and nutrient starvation responses has been done in lab strains which have been selected to be planktonic, making them easier to manipulate and assay in the lab. Second, MLP formation can often take days to manifest, as is evident in our experiments and other work (25), while most measurements for stress or starvation responses are taken a few minutes to a few hours after induction (100,110–113). Future experiments with longer timepoints, covering colony-level responses on the order of days (74), and accounting for cell-to-cell heterogeneity (e.g. single-cell RNA-seq (114–116), and strain-to-strain variability (e.g. within natural isolate libraries) will be fundamental for a more complete understanding of how yeast cells cope with environmental perturbations. In the slime mould *Dictyostelium discoideum,* a model organism with a facultative multicellular-like state, the transcriptomic landscape of the transition from unicellularity into multicellularity has been mapped at single-cell resolution (117). A similar experiment on planktonic *S. pombe* cells transitioning into flocculation might shed light on fundamental cellular decision-making processes and bet-hedging strategies (for example, in the case of cells that do not join flocs).

In this work, we generated valuable datasets that will form the basis of future mechanistic studies of MLP formation in *S. pombe*. Additionally, our work makes the first step towards understanding the natural diversity of MLP formation in fission yeast. Furthermore, we report novel players in MLP formation, some of which might represent pathways unique to *S. pombe*, and others which are conserved in other yeasts. Finally, our high-throughput assays of flocculation and surface adhesion are applicable to other microbes, and due to their high-throughput nature they could be used to uncover the diversity in MLP formation both within and across species. These assays can also be adopted for other large-scale experiments such as synthetic genetic arrays.

## Data availability and reproducibility

All collected data, performed analyses, and the sequence of the primers used have been deposited to https://github.com/BKover99/S.-Pombe-MLPs. Most analyses are available in a Jupyter notebook format (.ipynb). QTL analysis is available as an R script, while the haplotype calling pipeline is available as a bash script. The analysis tools used for our high-throughput assays can be accessed as a standalone package from https://github.com/BKover99/yeastmlp and can be installed from PyPI using the command “pip install yeastmlp”.

## Acknowledgements

We thank M. Lera-Ramirez and M.J. D’Angiolo for critical comments on the manuscript and S. Marguerat, D. Ellis, A. Garg, D. Jeffares, O. Hillson, B. Schwer, and S. Shuman for insightful discussions and sharing unpublished results.

## Notes

https://www.ebi.ac.uk/ena/browser/view/PRJEB69522

https://github.com/BKover99/S.-Pombe-MLPs

https://github.com/BKover99/yeastmlp

### Competing Interest Statement

The authors have declared no competing interest.

### Summary of Updates

Replaced figures with higher resolution images. Content unchanged.

