## supplemental_figures for "Genetic and environmental determinants of multicellular-like phenotypes in fission yeast"

### Supplementary Figure 1

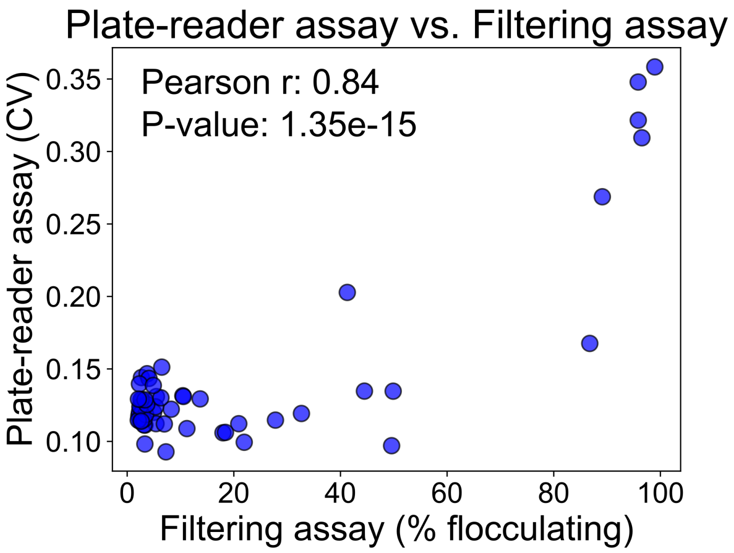


**Supplementary Figure 1**: Scatterplot comparison of flocculation for the JB50-JB759 segregant library by measuring each strain with a conventional filtering assay (x-axis, n=3) and our high-throughput plate-reader assay which measures the CV of OD600 within each well (y-axis, n=5) (Methods). Each point represents the mean flocculation values for a strain.

#### Supplementary Figure 2

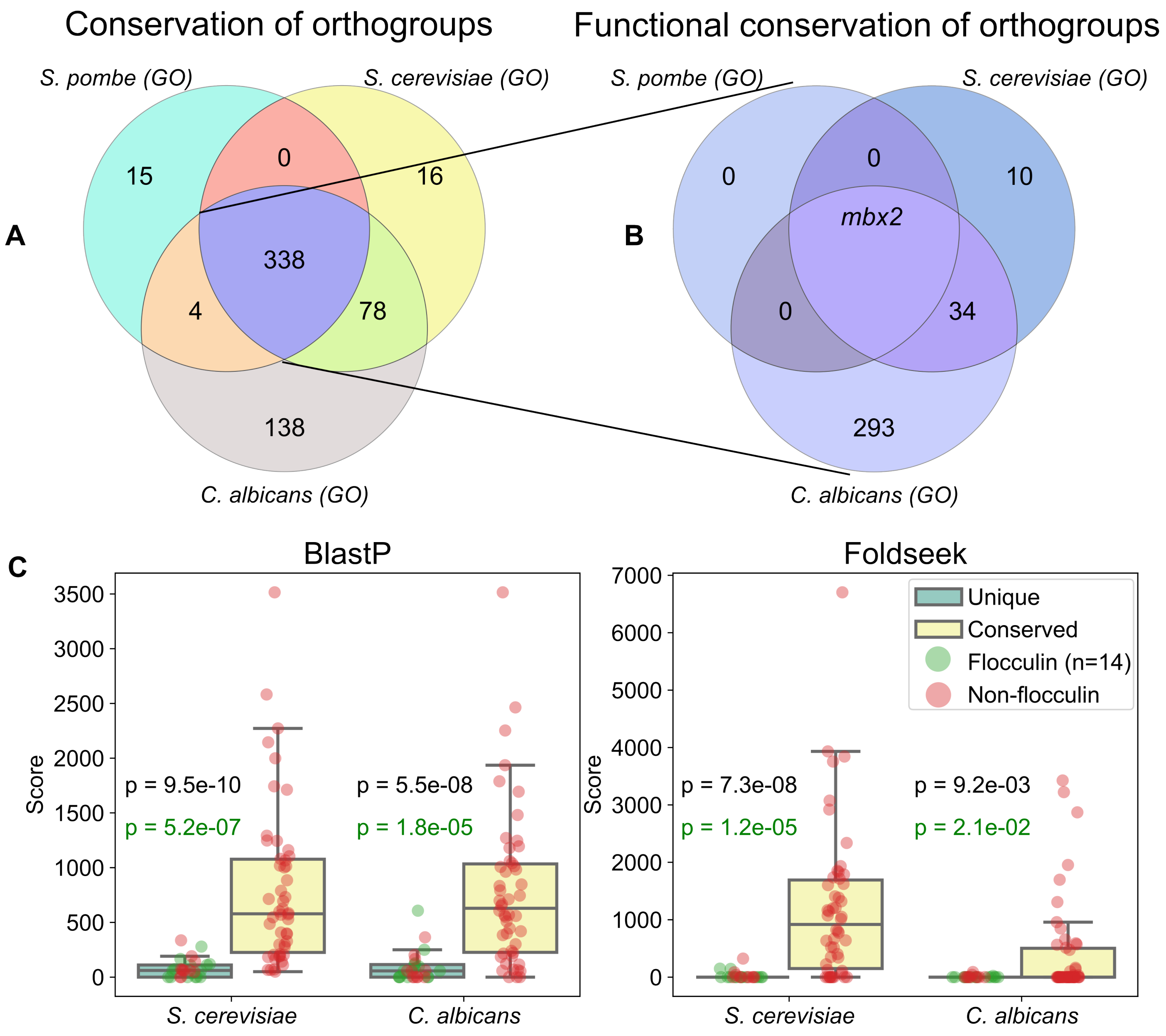


**Supplementary Figure 2**: Conservation of MLP-related orthogroups between S. pombe, S. cerevisiae and C. albicans. A: Venn diagram representing genetic and functional conservation of orthogroups between the three species using data only from GO-terms. B: Comparison of “unique” and “conserved” genes using BLAST-P and Foldseek, see Methods. Each dot represents a given gene/protein. P-values were derived from Mann-Whitney U tests. Black P-values were derived by comparing all “unique” and “conserved” genes. Green P-values were derived using the values of S. pombe flocculin genes/proteins only, rather than all “unique” MLP-related genes/proteins.

#### Supplementary Figure 3

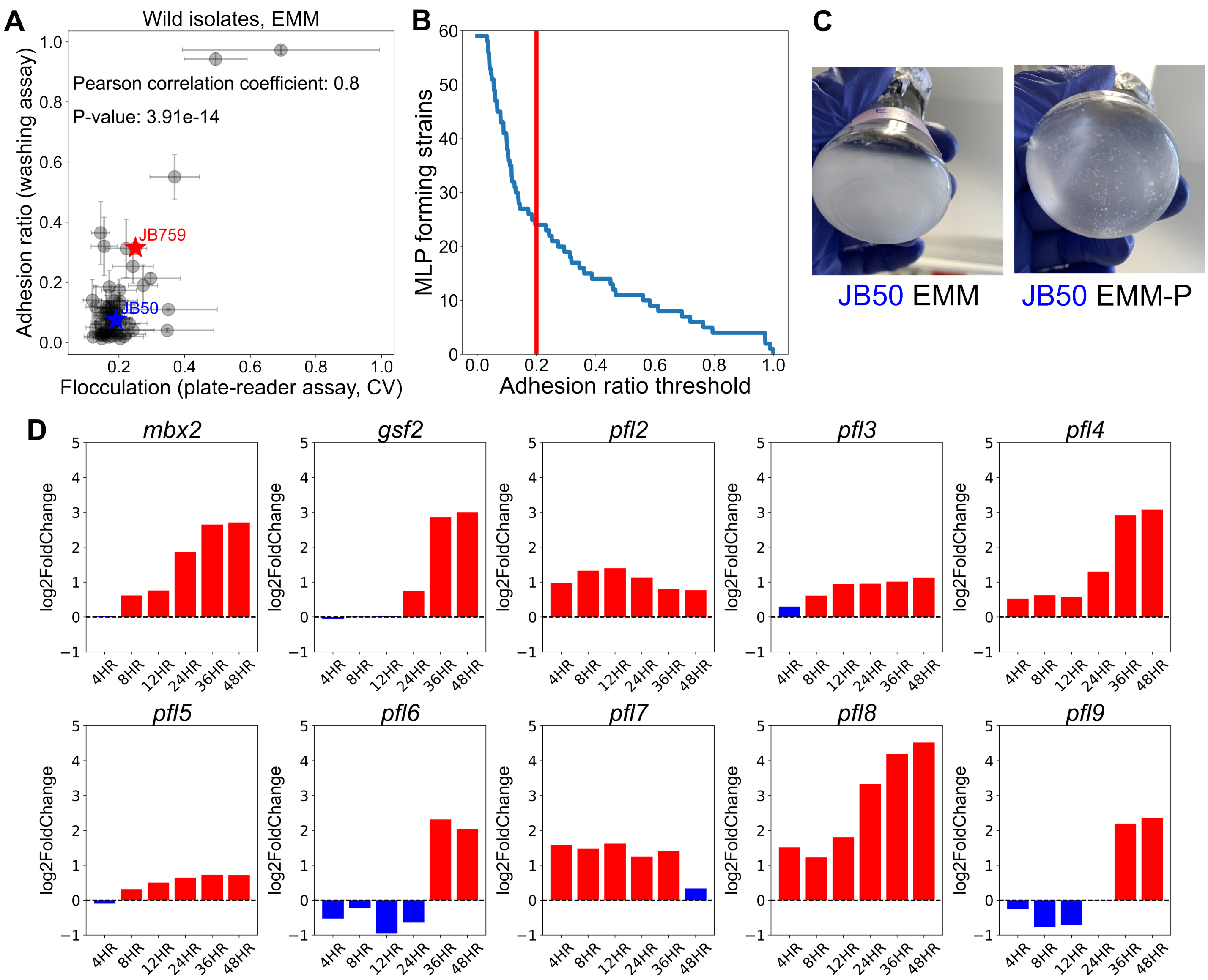


**Supplementary Figure 3**: Further analysis on adhesion phenotypes of the natural isolates across different conditions. A: Scatterplot comparison of flocculation and adhesion for the natural isolate library by measuring each strain with a high-throughput plate-reader assay (x-axis, n=11) and our high-throughput adhesion assay (y-axis, n=10). Each point represents the mean values for a strain with the standard error of the mean as error bars. Stars mark the our two strains of interest JB759 (red) and JB50 (blue). B: Elbow plot of the number of strains designated as "MLP-forming" in at least one condition (y-axis) given different cut-off thresholds (x-axis). Empirically we found that 0.2 sits in the elbow of the plot, suggesting that it is a reasonable cut-off. C: Phosphate starvation induces MLP formation in the lab strain JB50. Images of the JB50 strain growing in 25ml flasks with 10ml media of either EMM (left) or EMM-P (right). Strains were first grown up on YES plates and then transferred to liquid cultures. There are clearly visible flocs forming in EMM-P. E: Barplot of log2FC in mbx2 and flocculin expression at different timepoints during 2 days of phosphate starvation. Red bars represent significant log2FC values. Data taken from (74).

### Supplementary Figure 4

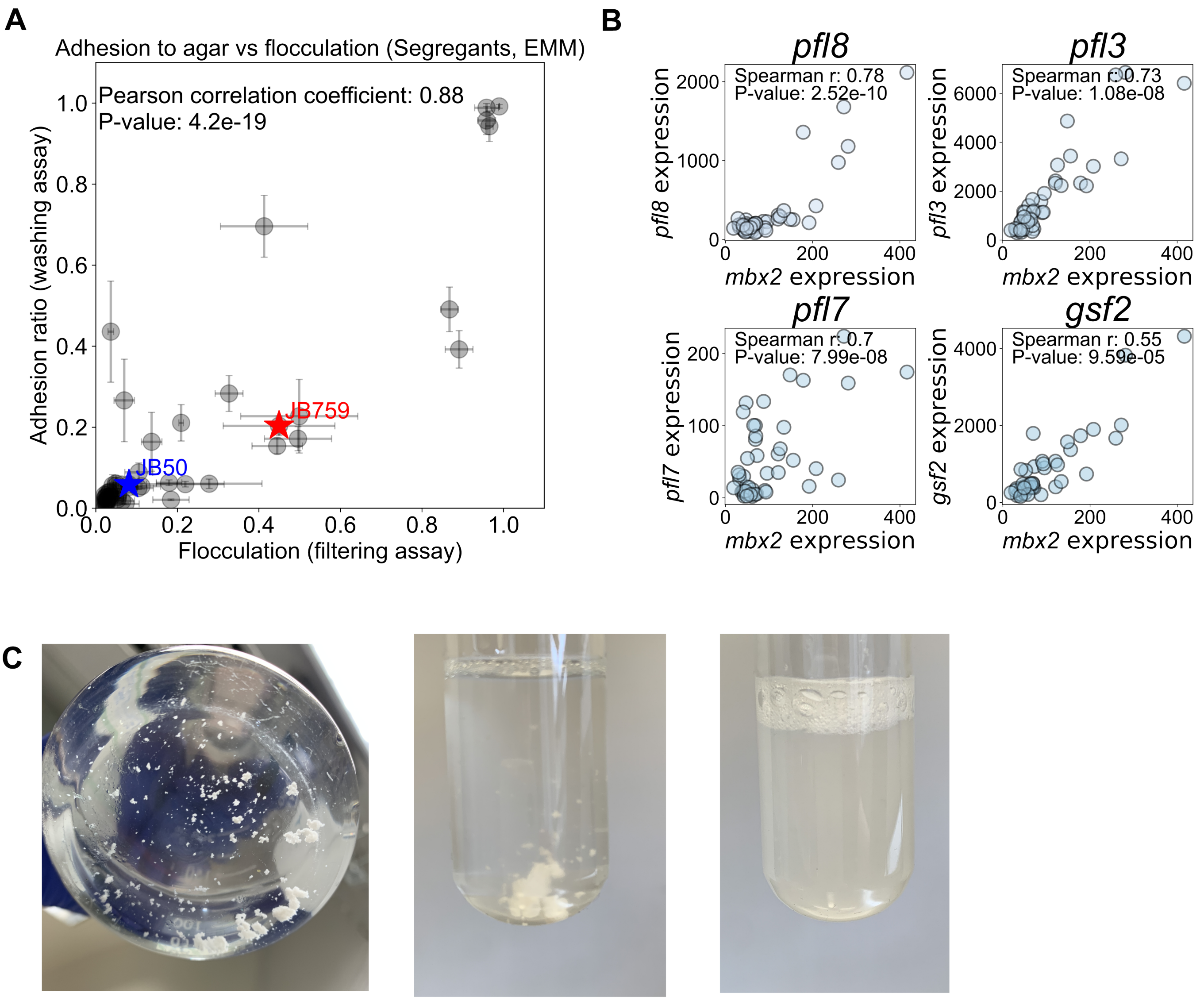


**Supplementary Figure 4**: Further analysis of the JB50-JB759 segregants and the role of mbx2. A: Scatterplot comparison of flocculation and adhesion for the JB50-JB759 library by measuring each strain with a filtering assay (x-axis, n=3) and our high-throughput adhesion assay (y-axis, n=10). Each point represents the mean values for a strain with the standard error of the mean as error bars. Stars mark the two parental strains JB759 (red) and JB50 (blue). There is a strong correlation between the two MLPs. B: Scatterplot of expression levels of four flocculin genes against mbx2 expression in the JB50-JB759 library (data from (46)), demonstrating a strong association. C: Results from the mbx2 overexpression strain on EMM. (Left, Middle) Images of flocculating mbx2OE strain in EMM. (Right) Images of mbx2OE strain after thiamine addition resulting in no flocculation.

### Supplementary Figure 5

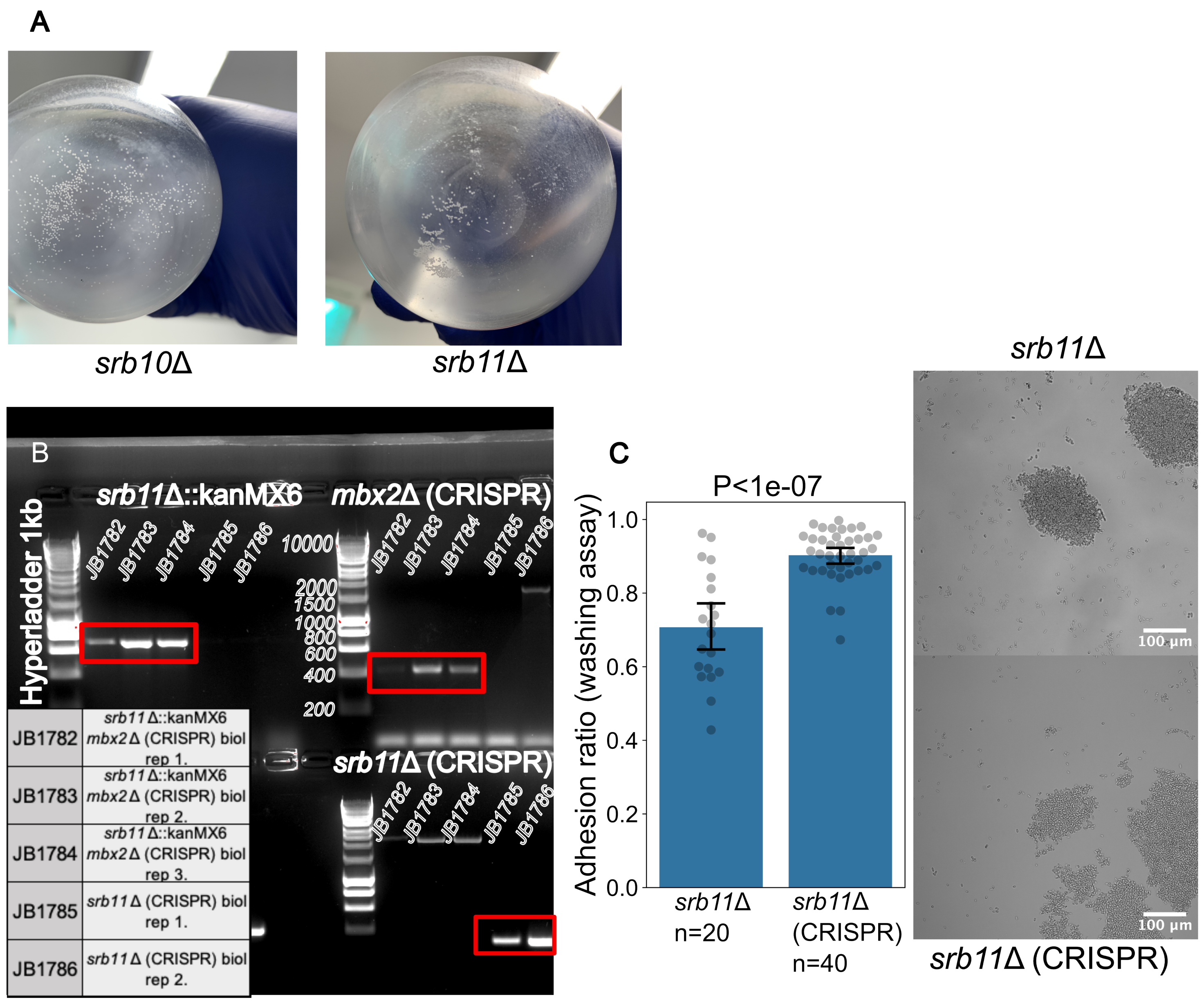


**Supplementary Figure 5**: The srb10Δ and srb11Δ strains show MLP formation. A: Images showing flocculating cultures of srb10 and srb11 in 250ml flasks grown in EMM. B: Validation of srb11Δ::Kan and CRISPR deletion strains used in the paper. Results from gel electrophoresis after PCR. Red rectangles mark the expected bands validating the strains. C: (Left) Barplot of adhesion values showing that the CRISPR deletion of srb11 phenocopies the Kan deletion of srb11 found in the deletion library. Additionally, we note that the CRISPR strain exhibits significantly stronger adhesion, likely because the deletion library strain has accumulated suppression mutations in other genes. Each dot is an independent measurement. Error bars represent the 95% confidence interval. (Right) Representative microscopy images of srb11Δ::Kan (top) and srb11Δ (CRISPR) (bottom) strains.

#### Supplementary Figure 6

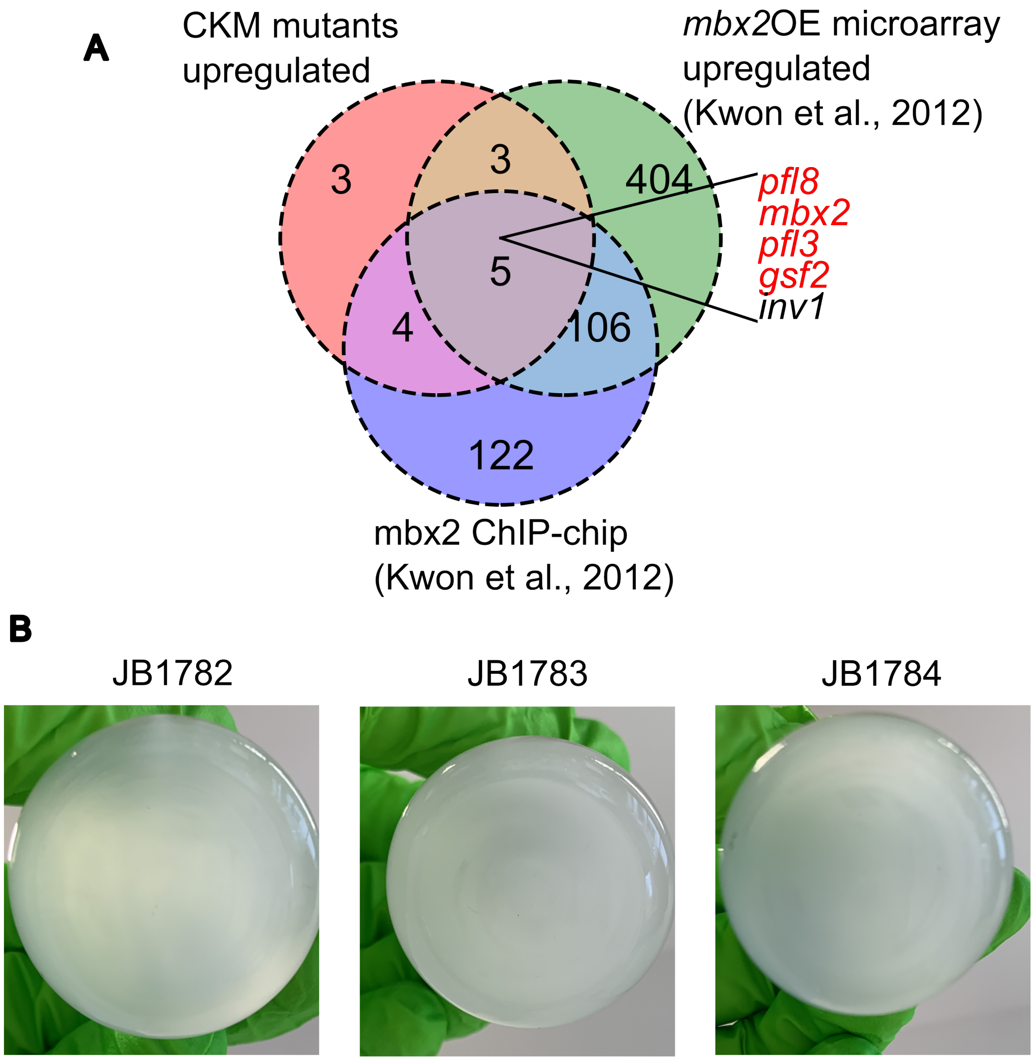


**Supplementary Figure 6**: Mbx2 drives MLP formation in the srb10Δ strain. A: Venn diagram showing the intersection of three gene sets: (i) Genes upregulated in CKM mutants (Fig 4E), meaning upregulated in the med12Δ and srb10Δ microarray data from (60) and in the srb11 truncated segregant RNA-seq data from (46), (ii) Genes upregulated after mbx2 overexpression obtained from (25) and (iii) Genes bound by Mbx2 obtained from (25). B: Images showing non-flocculating cultures of three biological replicates (JB1782, JB1783, JB1784) of srb11/mbx2 double delete strains.

### Supplementary Figure 7

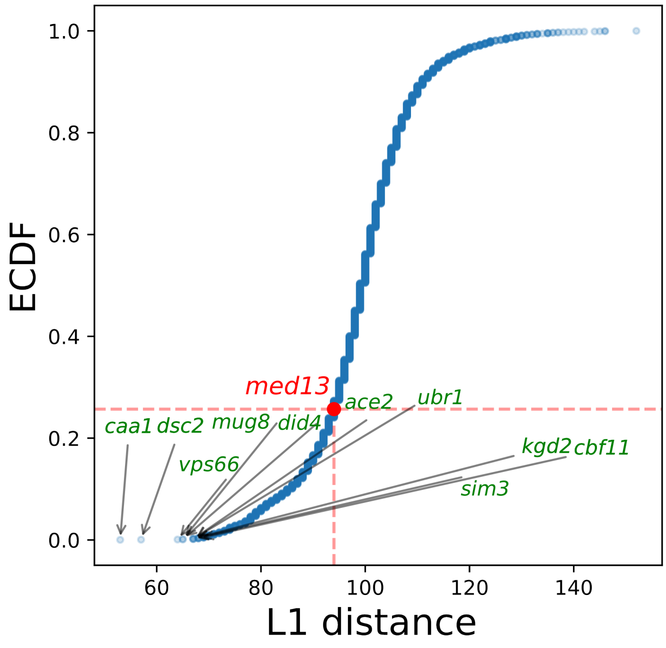


**Supplementary Figure 7**: In terms of the sensitivity/resistance phenotypes measured in (88), srb11Δ is not any more similar to med13Δ than to an average deletion strain. Empirical cumulative distribution function of L1 distances (sum of differences across phenotypes; -1,0,1 are sensitive, neutral and resistant). Top 10 deletion strains closest in their phenotypes to srb11Δ are highlighted on the plot.

### Supplementary Figure 8

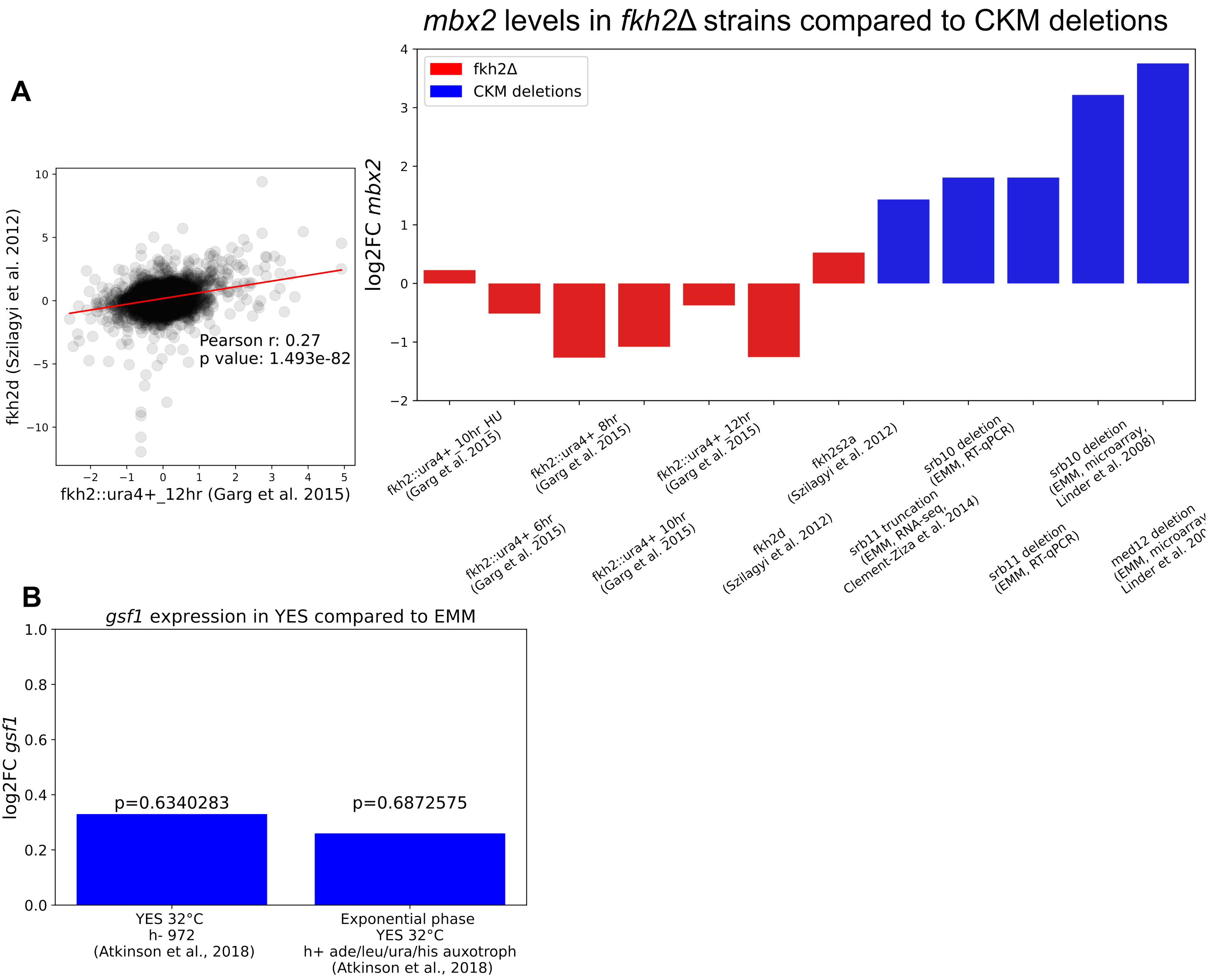


**Supplementary Figure 8**: Further analysis to deepen our understanding of the EMM-dependent upregulation of mbx2 in CKM deletions. Fkh2 is the only well-studied protein known to interact with the CKM in S. pombe, while gsf1 is the only known transcription factor to repress mbx2. A: Scatterplot of fold changes comparing two different data sources for transcriptomic changes upon fkh2 deletion. Each dot is a given gene in the dataset and the red line being a simple line of best fit. This plot concludes that it is fair to use the two data sources to draw joint conclusions from them. B: Barplot showing log2 fold-changes in mbx2 expression levels upon fkh2 deletion from (79) and (89) compared with expected fold changes from med12Δ and srb10Δ microarray data from (60) and in the srb11 truncated segregant RNA-seq data from (46) and our RT-qPCR data collected in this work. C: Barplot showing log2 fold-changes in gsf1 expression in YES compared to EMM. There is no significant upregulation in either of the two replicates. Data and P-values were taken from (90).

### Supplementary Figure 9

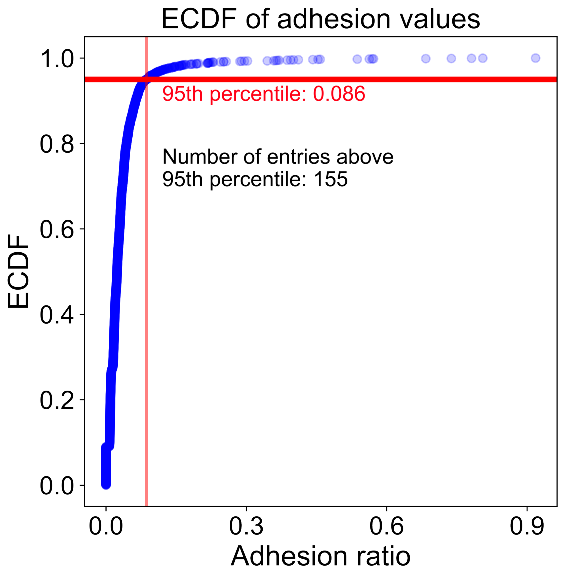


**Supplementary Figure 9**: Empirical cumulative distribution function (ECDF) of adhesion ratios from the deletion library screen, with red lines showing the cut-off for the 95th percentile. Strains above the cut-off were used for enrichment analysis.

### Supplementary Figure 10

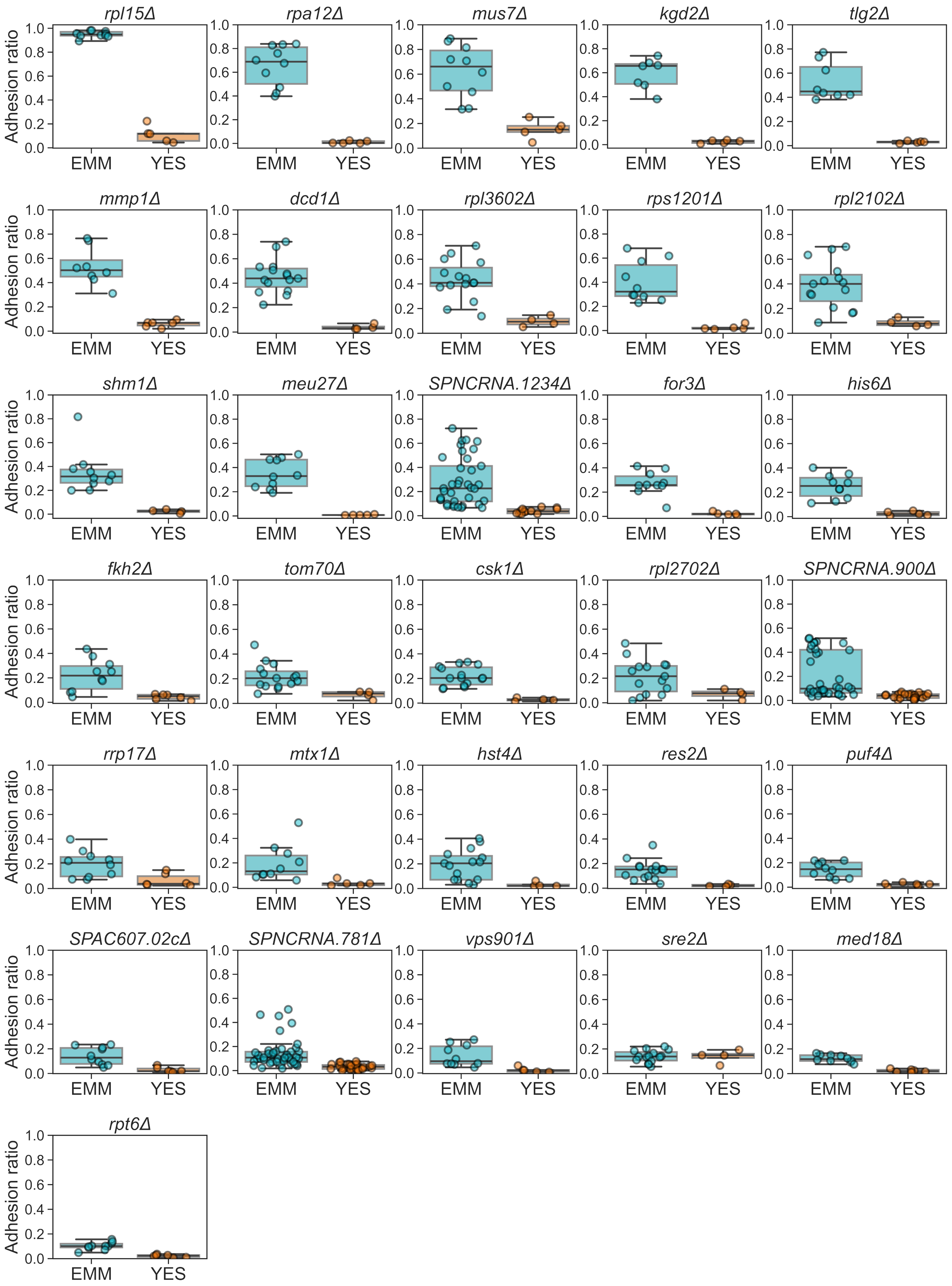

**Supplementary Figure 10**: Strip-boxplots of adhesion ratios obtained with the washing assay for the 31 high-confidence hits on EMM (light blue) vs YES (orange). Each dot is an independent observation. Adhesion was generally specific to EMM, except in the case of sre2Δ.

### Supplementary Figure 11

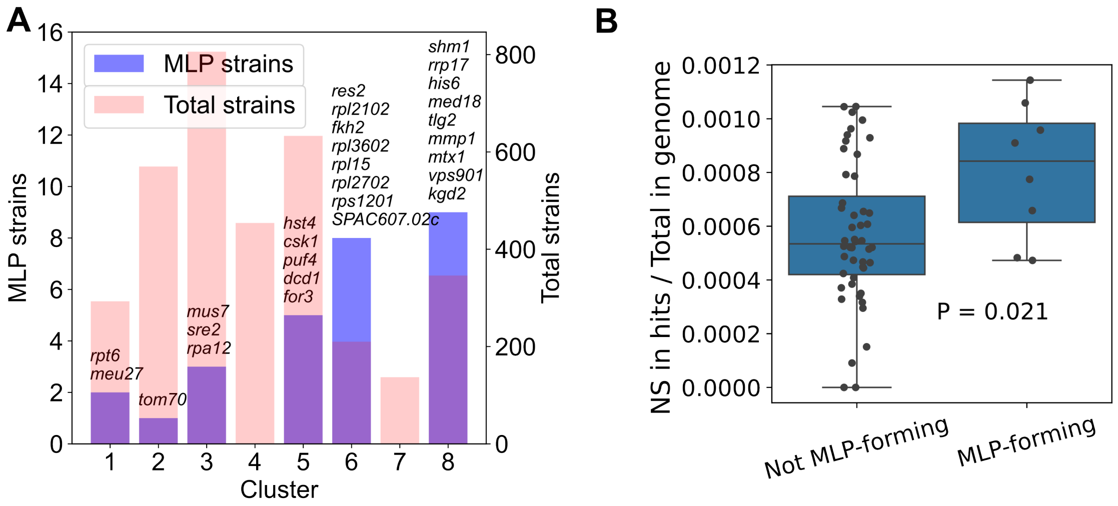


**Supplementary Figure 11**: A: Barplot showing the number of MLP-forming (red) and total (blue) strains in each broad phenotypic cluster of deletion strains identified in (88). B: Strip-boxplot showing non-synonymous SNPs in the 31 hits from the deletion screen in natural isolates that exhibit MLPs in EMM vs natural isolates that do not exhibit MLPs in EMM. The total number of non-synonymous SNPs in the 31 genes was summed and divided by the total number of mutations (relative to the reference strain) for that strain. Each dot marks a natural isolate. P-value was calculated using a permutation-based T-test with 100 000 permutations.

#### Supplementary Figure Legends

**Supplementary figure 1:** Scatterplot comparison of flocculation for the JB50-JB759 segregant library by measuring each strain with a conventional filtering assay (x-axis, n=3) and our high-throughput plate-reader assay which measures the CV of OD600 within each well (y-axis, n=5) (Methods). Each point represents the mean flocculation values for a strain.

**Supplementary figure 2: Conservation of MLP-related orthogroups between *S. pombe, S. cerevisiae* and *C. albicans*.** A: Venn diagram representing genetic and functional conservation of orthogroups between the three species using data only from GO-terms. B: Comparison of “unique” and “conserved” genes using BLAST-P and Foldseek, see Methods. Each dot represents a given gene/protein. P-values were derived from Mann-Whitney U tests. Black P-values were derived by comparing all “unique” and “conserved” genes. Green P-values were derived using the values of *S. pombe* flocculin genes/proteins only, rather than all “unique” MLP-related genes/proteins.

**Supplementary figure 3: Further analysis on adhesion phenotypes of the natural isolates across different conditions.** A: Scatterplot comparison of flocculation and adhesion for the natural isolate library by measuring each strain with a high-throughput plate-reader assay (x-axis, n=11) and our high-throughput adhesion assay (y-axis, n=10). Each point represents the mean values for a strain with the standard error of the mean as error bars. Stars mark the our two strains of interest JB759 (red) and JB50 (blue). B: Elbow plot of the number of strains designated as “MLP-forming” in at least one condition (y-axis) given different cut-off thresholds (x-axis). Empirically we found that 0.2 sits in the elbow of the plot, suggesting that it is a reasonable cut-off. C: Phosphate starvation induces MLP formation in the lab strain JB50. Images of the JB50 strain growing in 25ml flasks with 10ml media of either EMM (left) or EMM-P (right). Strains were first grown up on YES plates and then transferred to liquid cultures. There are clearly visible flocs forming in EMM-P. E: Barplot of log2FC in *mbx2* and flocculin expression at different timepoints during 2 days of phosphate starvation. Red bars represent significant log2FC values. Data taken from (74).

**Supplementary figure 4: Further analysis of the JB50-JB759 segregants and the role of *mbx2*.** A: Scatterplot comparison of flocculation and adhesion for the JB50-JB759 library by measuring each strain with a filtering assay (x-axis, n=3) and our high-throughput adhesion assay (y-axis, n=10). Each point represents the mean values for a strain with the standard error of the mean as error bars. Stars mark the two parental strains JB759 (red) and JB50 (blue). There is a strong correlation between the two MLPs. B: Scatterplot of expression levels of four flocculin genes against *mbx2* expression in the JB50-JB759 library (data from (46)), demonstrating a strong association. C: Results from the *mbx2* overexpression strain on EMM. (Left, Middle) Images of flocculating mbx2OE strain in EMM. (Right) Images of *mbx2OE* strain after thiamine addition resulting in no flocculation.

**Supplementary figure 5: The *srb10*Δ and *srb11*Δ strains show MLP formation.** A: Images showing flocculating cultures of *srb10* and *srb11* in 250ml flasks grown in EMM. B: Validation of *srb11Δ::Kan* and CRISPR deletion strains used in the paper. Results from gel electrophoresis after PCR. Red rectangles mark the expected bands validating the strains. C: (Left) Barplot of adhesion values showing that the CRISPR deletion of *srb11* phenocopies the *Kan* deletion of *srb11* found in the deletion library. Additionally, we note that the CRISPR strain exhibits significantly stronger adhesion, likely because the deletion library strain has accumulated suppression mutations in other genes. Each dot is an independent measurement. Error bars represent the 95% confidence interval. (Right) Representative microscopy images of *srb11*Δ::*Kan* (top) and *srb11*Δ (CRISPR) (bottom) strains.

**Supplementary figure 6: Mbx2 drives MLP formation in the *srb10*Δ strain.** A: Venn diagram showing the intersection of three gene sets: (i) Genes upregulated in CKM mutants (Fig 4E), meaning upregulated in the *med12*Δ and *srb10*Δ microarray data from (60) and in the *srb11* truncated segregant RNA-seq data from (46), (ii) Genes upregulated after *mbx2* overexpression obtained from (25) and (iii) Genes bound by Mbx2 obtained from (25). B: Images showing non-flocculating cultures of three biological replicates (JB1782, JB1783, JB1784) of *srb11/mbx2* double delete strains.

**Supplementary figure 7:** In terms of the sensitivity/resistance phenotypes measured in (88), *srb11*Δ is not any more similar to *med13*Δ than to an average deletion strain. Empirical cumulative distribution function of L1 distances (sum of differences across phenotypes; -1,0,1 are sensitive, neutral and resistant). Top 10 deletion strains closest in their phenotypes to *srb11*Δ are highlighted on the plot.

**Supplementary figure 8: Further analysis to deepen our understanding of the EMM-dependent upregulation of *mbx2* in CKM deletions.** Fkh2 is the only well-studied protein known to interact with the CKM in *S. pombe*, while *gsf1* is the only known transcription factor to repress *mbx2*. A: Scatterplot of fold changes comparing two different data sources for transcriptomic changes upon *fkh2* deletion. Each dot is a given gene in the dataset and the red line being a simple line of best fit. This plot concludes that it is fair to use the two data sources to draw joint conclusions from them. B: Barplot showing log2 fold-changes in *mbx2* expression levels upon *fkh2* deletion from (79) and (89) compared with expected fold changes from *med12*Δ and *srb10*Δ microarray data from (60) and in the *srb11* truncated segregant RNA-seq data from (46) and our RT-qPCR data collected in this work. C: Barplot showing log2 fold-changes in *gsf1* expression in YES compared to EMM. There is no significant upregulation in either of the two replicates. Data and P-values were taken from (90).

**Supplementary figure 9:** Empirical cumulative distribution function (ECDF) of adhesion ratios from the deletion library screen, with red lines showing the cut-off for the 95th percentile. Strains above the cut-off were used for enrichment analysis.

**Supplementary figure 10:** Strip-boxplots of adhesion ratios obtained with the washing assay for the 31 high-confidence hits on EMM (light blue) vs YES (orange). Each dot is an independent observation. Adhesion was generally specific to EMM, except in the case of *sre2Δ.*

**Supplementary figure 11:** A: Barplot showing the number of MLP-forming (red) and total (blue) strains in each broad phenotypic cluster of deletion strains identified in (88). B: Strip-boxplot showing non-synonymous SNPs in the 31 hits from the deletion screen in natural isolates that exhibit MLPs in EMM vs natural isolates that do not exhibit MLPs in EMM. The total number of non-synonymous SNPs in the 31 genes was summed and divided by the total number of mutations (relative to the reference strain) for that strain. Each dot marks a natural isolate. P-value was calculated using a permutation-based T-test with 100 000 permutations.
